# eIF4E assembly into *C. elegans* germ granules is essential for its repressive function

**DOI:** 10.1101/2025.09.24.678427

**Authors:** Carmen Herrera Sandoval, Christopher Borchers, Boyoon Yang, Becky Boyd, Heather A. Hundley, Scott T. Aoki

## Abstract

Metazoan germ cells form intracellular germ granules, cytoplasmic RNA-protein condensates that contain a variety of RNAs and proteins essential for germline identity, maintenance, and fertility. P granules are a type of *C. elegans* germ granule proposed to be sites of mRNA repression. Proper P granule assembly is dependent on PGL-1 and its granule-forming protein relatives. Numerous RNA-binding proteins localize to P granules, like the eIF4E mRNA cap binding homolog, IFE-1. IFE-1 directly interacts with PGL-1 in vivo and in vitro. The molecular function of P granules remains enigmatic. Here, PGL-1 was molecularly dissected in vivo to determine protein regions required for P granule assembly, binding partner recruitment, and germ cell development. A specific region in the PGL-1 C-terminus was necessary and sufficient for IFE-1 recruitment to P granules and for fertility. IFE-1 RNA targets were identified, and reporters of top gene targets were repressed in the adult germline. This repression was dependent on PGL-1 and its IFE-1 binding peptide. These findings provide evidence that IFE-1 and P granules are a factor and site of mRNA repression, respectively. This repression required IFE-1 assembly into P granules, supporting the model that RNA-protein condensate assembly is necessary for its biological and biochemical functions.

## Introduction

Effective cellular function depends on organizing macromolecules into compartments to facilitate biochemistry. Biomolecular condensates are non-membrane bound compartments that include nucleoli, stress granules, and metazoan germ granules [1]. They assemble by liquid-liquid phase separation [1], a biophysical property driven by favorable intermolecular interactions that exceed those with the surrounding cytoplasm. Biomolecular condensation depends on proteins and other molecules with multiple interaction domains and/or intrinsically disordered regions (IDRs) [1–3]. Although many biomolecular condensates have been identified, their mechanisms of formation and necessity of condensation for their biological functions remain largely elusive.

Germ granules are cytoplasmic RNA-protein condensates found in the metazoan germ cells [4, 5]. Although the specific composition of germ granules differs across species, they often recruit mRNA regulatory factors and regulate mRNAs post-transcriptionally to influence development and protect germ cells. In *C. elegans,* P granules are a type of germ granule required for proper germ cell development [4, 5]. They localize to the nuclear periphery for most of adult germ cell development [6, 7] and behave as phase-separated, liquid condensates [8]. By electron microscopy, P granules are electron-dense due to their high RNA content [9]. Numerous RNA-binding and regulatory proteins localize to P granules [5] such as IFE-1, a germline specific eIF4E homolog involved in sperm regulation [10, 11] and oocyte maturation [12, 13]. eIF4E recognizes the 7-methylguanosine (m^7^G) 5’ cap on mRNAs and interacts with other proteins to regulate mRNA localization, storage, turnover, and both translational activation and repression [14]. The condensate assembly protein PGL-1 mediates the recruitment of IFE-1 to P granules, observed both in vivo [10, 11] and in vitro [6, 10]. During spermatogensis, it is speculated that IFE-1 is released from P granules into the cytoplasm where it may facilitate spermatogenesis [10]. Indeed, *ife-1* null mutants are sterile at high temperatures primarily due to defects in sperm function [10–12]. Others suggested that IFE-1 acts as a repressor of its target mRNAs when bound to PGL-1 [15], but the specific interaction between IFE-1 and PGL-1, IFE-1 target mRNA repression, and necessity of granule assembly for IFE-1 function remain undiscovered.

In *C. elegans*, proper P granule assembly is dependent on the PGL family of granule-forming proteins, which include PGL-1 and PGL-3 [6, 7]. These proteins have similar sequences and functional redundance [6, 16], although only PGL-1 binds to IFE-1 [10]. Both PGL-1 and PGL-3 are required for proper germ cell development [6]. Simultaneous genetic removal of these proteins leads to abnormal expression of spermatogenic and somatic mRNAs [17], resulting in sterility [6, 7]. These two PGL proteins can be divided into three distinct regions: (1) the N-terminal domain (Nt), (2) the dimerization domain (DD), and (3) the C-terminal region (Ct) with RGG repeats (RGG) **(Fig. 1A)**. The structure of the Nt and DD domains have been resolved [18, 19], and dimerization of the Nt domain has been shown to be critical for P granule assembly [18, 20, 21]. Functional studies using a lambda peptide-stem loop system demonstrated that artificially tethering reporter mRNAs to PGL-1 recruited them to P granules and repressed their translation [18]. This repression required Nt domain dimerization, supporting the idea that condensate formation is necessary for mRNA regulation. However, this experimental system has two key limitations. First, the lambda peptide insertion is artificial and may not reflect endogenous PGL-1 interactions. Second, homozygous expression of lambda-tagged PGL-1 caused sterility [18], requiring experiments to be performed in heterozygous animals, further limiting physiological relevance. Moreover, Nt domain dimerization may contribute to biochemical functions beyond condensate assembly. Therefore, additional studies are needed to directly test whether P granules are required for endogenous mRNA repression.

**Figure 1.**
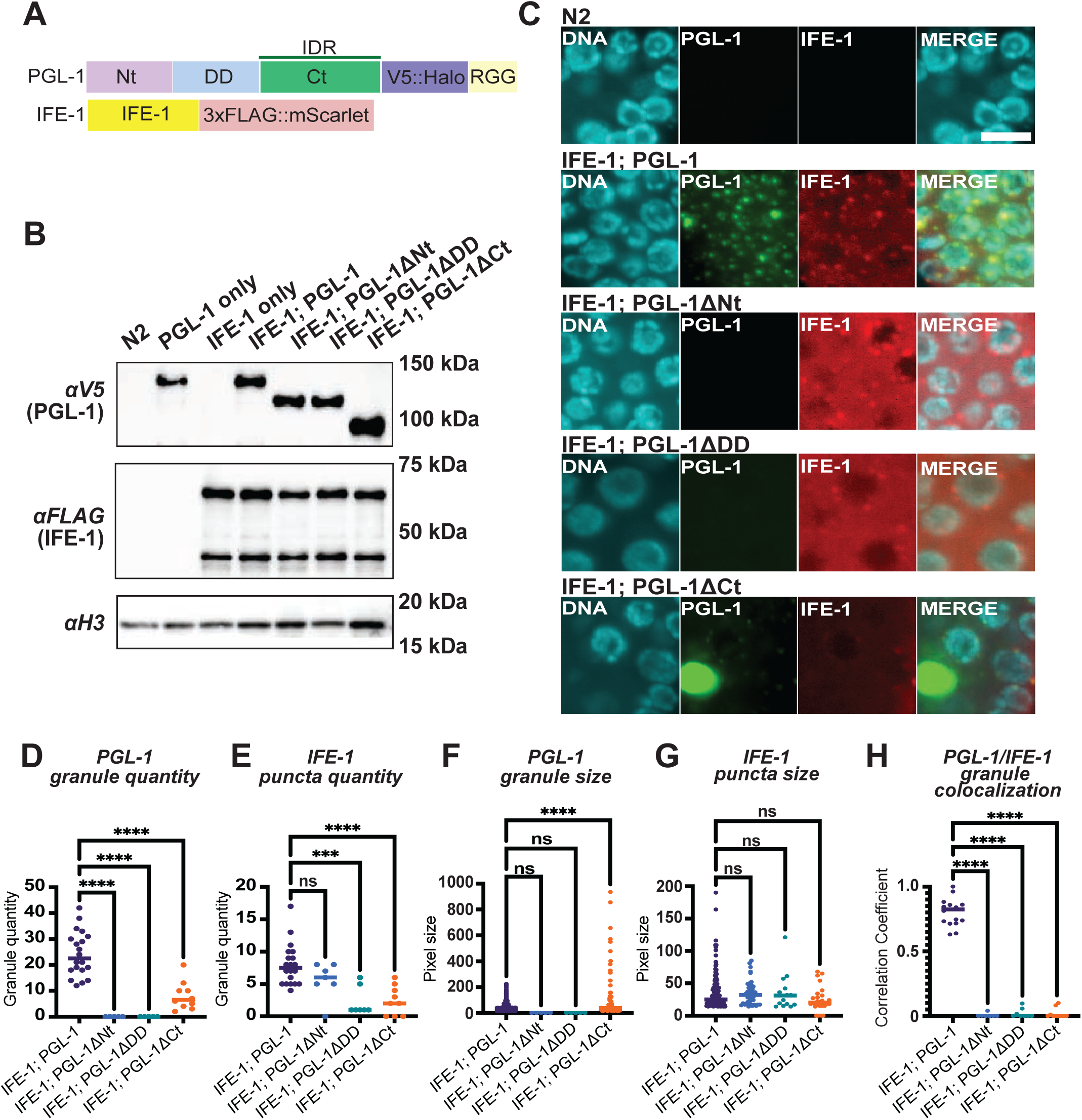
Deletion of the PGL-1 C-terminal region disrupts normal granule formation. (A) Linear diagram of *C. elegans* PGL-1 and IFE-1. Nt, N-terminal domain (light purple); DD, dimerization domain, (light blue); Ct, C-terminus region (green); V5 and Halo tag (blue-magenta); RGG repeats (light yellow); IFE-1 (yellow); 3xFLAG and mScarlet tag (light red); intrinsically disordered region, IDR. Diagram not to scale. (B) Immunoblot probing PGL-1 and IFE-1 protein expression. Adult hermaphrodite lysates were separated by SDS-PAGE and probed with V5 to detect PGL-1, FLAG to detect IFE-1, and histone H3 (H3) antibodies. Strains expressed IFE-1::3xFLAG::mScarlet (IFE-1) with PGL-1::V5::Halo (PGL-1), PGL-1ΔNt::V5::Halo (PGL-1ΔNt), PGL-1ΔDD::V5::Halo (PGL-1ΔDD), PGL-1ΔCt::V5::Halo (PGL-1ΔCt). N2 wildtype, PGL-1 only, and IFE-1 only served as negative controls. (**C**) Confocal microscopy of adult hermaphrodite germlines. Strains imaged carried wild-type or mutant PGL-1 with 3xFLAG::mScarlet-tagged IFE-1. DNA, DAPI (cyan); PGL-1, Halo Oregon Green (green); IFE-1, αFLAG (red); and a merged panel. Representative images are 10-layer z-stack maximum intensity projections. Scale bar, white, 5 μm. (**D-G**) Confocal image analyses of PGL-1 granule and IFE-1 puncta. PGL-1 granule (D) and IFE-1 puncta (E) quantification. PGL-1 granule (F) and IFE-1 puncta (G) size quantification. (**H**) Correlation of IFE-1 puncta colocalizing with PGL-1 granules. Ordinary one-way ANOVA statistical test was used to compare data. ns, not significant; ***, p-value 0.001; ****, p-value <0.0001.

To investigate the molecular mechanisms driving P granule assembly, genetic dissection of the PGL-1 protein was performed in vivo. Deletion of regions in PGL-1 Ct resulted in abnormal P granule formation and IFE-1 localization. IFE-1-associated RNAs were identified, and high confidence target mRNA transcripts were repressed in the adult germline. This repression required IFE-1 recruitment to P granules. Together, these findings demonstrate that proper assembly of IFE-1 into P granules is necessary for its role in mRNA repression. The work advances our understanding of how a liquid RNA-protein condensate and germ granule assembly contributes to post-transcriptional gene regulation.

## Results

### The PGL-1 C-terminus influences P granule assembly

*C. elegans* PGL-1 is critical for proper P granule assembly [6, 7]. Prior studies identified multiple dimerization domains in PGL-1 [18, 19], with at least one required for P granule formation [18]. The C-terminal region of PGL-1 contains RGG repeats [7, 22], is predicted to be intrinsically disordered [19], and has patches of sequence conservation that suggest a potential role in recruiting or organizing proteins within granules **(Fig. S1A)**. IFE-1 is a germline-specific eIF4E homolog involved in sperm regulation [10, 11, 13, 15] and oocyte maturation [12]. IFE-1 binds to the m^7^G 5’ cap of mRNAs [10, 23], localizes to P granules [10], and directly interacts with PGL-1 in vivo [10, 11] and in vitro [6, 10]. The specific PGL-1 protein regions required for IFE-1 interaction are unknown.

A first goal was to determine the specific contributions of each PGL-1 region for P granule assembly and IFE-1 recruitment. CRISPR/Cas-9 gene editing was used to insert a V5 epitope [24] and HaloTag (Halo, [25]) in *pgl-1* and a 3xFLAG tagged mScarlet fluorescent protein in *ife-1* for straightforward imaging **(Fig. 1A)**. These tagged proteins are referred to as PGL-1 and IFE-1 hereafter for clarity. The N-terminal domain (Nt), Dimerization Domain (DD), and C-terminal region (Ct) of *pgl-1* **(Fig. 1A)** were deleted to test their involvement in assembly. Protein expression of all mutants was confirmed via immunoblots **(Fig. 1B).** Confocal microscopy assessed P granule assembly of the *pgl-1* deletion mutants. Wildtype N2 served as a negative control, and IFE-1; PGL-1 animals served as a positive control **(Fig. 1C, S1B)**. The positive control animals showed robust PGL-1 granule and IFE-1 puncta formation **(Fig. 1C-G)** that colocalized with each other **(Fig. 1H)**, as expected [6, 10]. In contrast, the removal of the PGL-1 Nt or DD domains disrupted P granule assembly **(Fig. 1C)**, both in PGL-1 granule quantity **(Fig. 1D)** and PGL-1 colocalization with IFE-1 **(Fig. 1H)**. Deletion of the PGL-1 Ct region also caused a significant reduction in the number of PGL-1 granules and IFE-1 puncta **(Fig. 1C-E)** and affected IFE-1 colocalization with PGL-1 **(Fig. 1H)**. Surprisingly, the few remaining P granules were incredibly enlarged, with a few as large as germ cell nuclei **(Fig. 1C,F).** IFE-1 puncta numbers were decreased in both the DD domain and Ct region deletion mutants **(Fig. 1C,E)** but the overall puncta size were similar in all strains **(Fig. 1C,G).** Of note, removal of the PGL-1 Ct region did not cause enlargement of the IFE-1 puncta **(Fig. 1C,G)**, implying abnormal IFE-1 recruitment. In sum, the imaging results suggested a requirement of the PGL-1 Ct region for proper PGL-1 granule assembly and IFE-1 co-factor recruitment.

Fertility assays were used to determine whether the PGL-1 deletions affected germ cell development. While *pgl-1 null (bn101)* mutants propagate under standard 20°C conditions **(Fig. 2A, S1C)**, they exhibit a dramatic 25°C temperature-sensitive fertility defect [7] **(Fig. 2B, S1D).** Fertility defects with PGL-1 mutants may thus be attributable to a loss in PGL-1 protein function. At 20°C, both wildtype N2 and Halo-tagged PGL-1 were fully fertile **(Fig. 2A, S1C)**, while removal of PGL-1 Nt, DD, and Ct regions caused decreased fertility **(Fig. 2A).** At 25°C, wildtype N2 remained fertile, *pgl-1 (bn101) null* worms were sterile, and Halo-tagged PGL-1 showed a modest decrease in fertility **(Fig. 2B, S1D)**, a result consistent with the effect of other large protein tags added to PGL-1 [18]. Others noted *ife-1* null animals to be completely sterile at 25°C [12]. The fertility observed in these strains carrying a tagged *ife-1* allele **(Fig. 2B, S1D)** suggested that the added tag did not grossly affect protein function. In contrast, all three PGL-1 deletion mutants exhibited near complete sterility at 25°C **(Fig. 2B)**. Collectively, these results indicate that the PGL-1 Nt, DD, and Ct regions contribute to both P granule assembly and its biological function in germ cell development. Although functions have previously been assigned to the Nt and DD domains [18, 19], the biochemical role of the PGL-1 Ct region was unknown.

**Figure 2.**
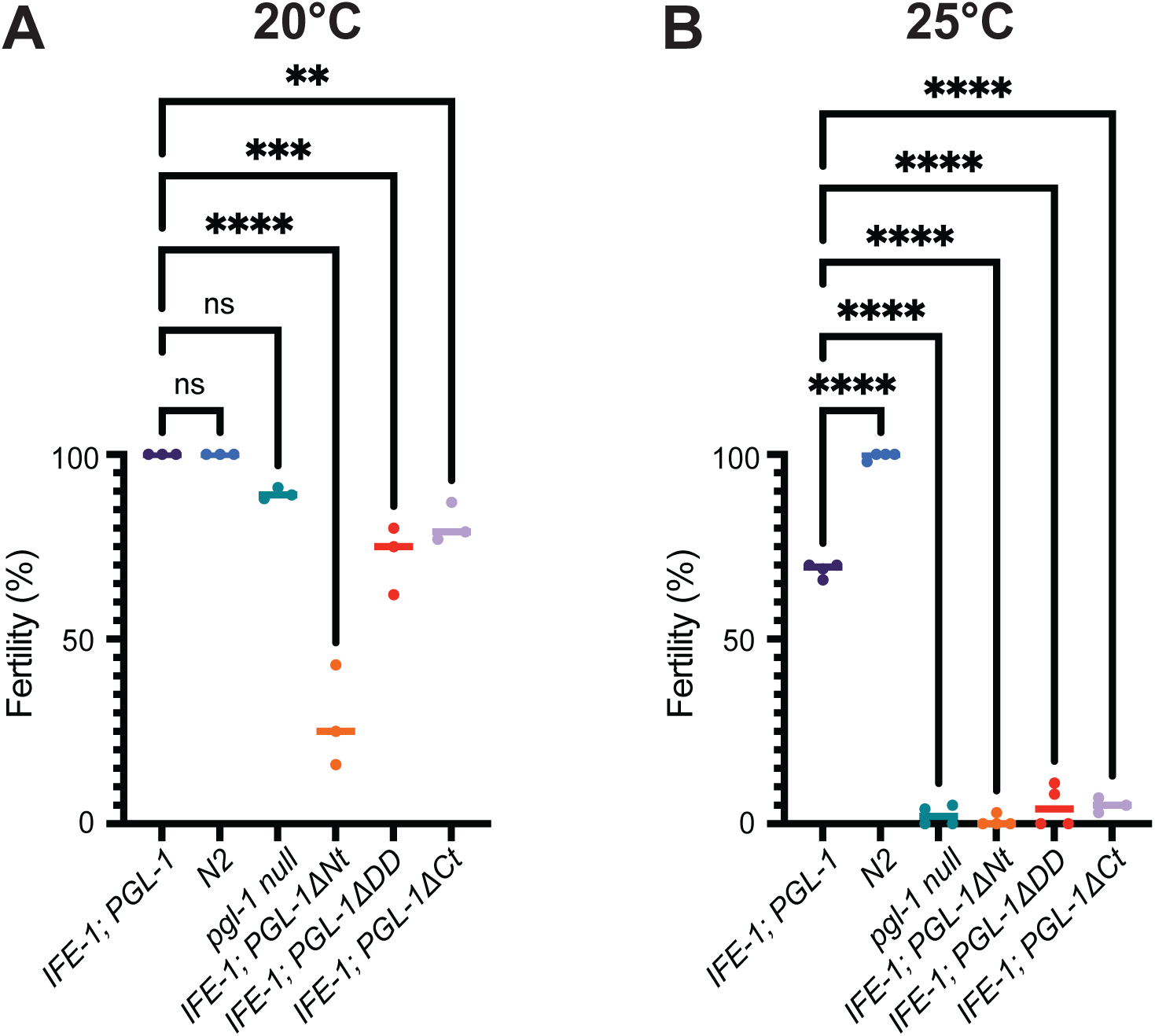
PGL-1 deletion mutants exhibit fertility defects. Fertility of Halo-tagged PGL-1 wildtype and mutant hermaphrodites. Worms were plated individually, propagated at (**A**) 20°C or (**B**) 25°C, and scored for presence of larval progeny after 4 days. Results reported as the percentage of fertile animals relative to the total number tested in each experiment. Strains expressed IFE-1::3xFLAG::mScarlet (IFE-1) with PGL-1::V5::Halo (PGL-1), PGL-1ΔNt::V5::Halo (PGL-1ΔNt), PGL-1ΔDD::V5::Halo (PGL-1ΔDD), PGL-1ΔCt::V5::Halo (PGL-1ΔCt). The *pgl-1 null* mutant strain served as an assay control for infertility, while N2 served as a wildtype control. At 25 °C, fertility is significantly decreased in all *pgl-1* mutant strains. *n =* 3 independent experiments. *, p-value <0.05; **, p-value <0.01; ****, p-value <0.0001.

### A PGL-1 C-terminal region is required for IFE-1 recruitment and proper germ cell development

Yeast-two-hybrid molecular genetics was used to screen for a potential function for the PGL-1 Ct. PGL-1 Ct was used as bait and screened with a library of *C. elegans* protein coding regions to identify potential protein-binding partners (see **Methods**). Three protein candidates were identified as high-confidence interactors: M01E5.4, LYS-7, and IFE-1. LYS-7 is a secreted lysozyme whose expression is induced upon pathogen infection [26]. Because LYS-7 is secreted, it is unlikely to function as a cytoplasmic P granule interactor. M01E5.4 is a human C2orf68 homolog [27] and poorly characterized in worms. CRISPR/Cas9 was used to insert a 3xFLAG::mScarlet tag in the endogenous *M01E5.4* locus **(Fig. S2A).** Despite confirming tagged M01E5.4 expression by immunoblot **(Fig. S2B)**, confocal imaging did not reveal germline protein expression **(Fig. S2C)**, implying that M01E5.4 does not interact with PGL-1 in the adult germline. IFE-1 is an established PGL-1 interactor [6, 10]. Given that IFE-1 puncta formation is disrupted in PGL-1 Ct deletion mutants **(Fig. 1C,E,H)**, a peptide region within PGL-1 Ct may directly bind IFE-1.

The PGL-1 Ct region was further dissected in vivo to identify protein regions responsible for P granule assembly, IFE-1 interaction, and germ cell development. PGL Ct protein alignments showed patches of conservation **(Fig. S1A)**. The Ct region was divided further into 3 roughly equal parts with boundaries in non-conserved regions, referred PGL-1 CtA, CtB, and CtC **(Fig. S1A)**. CRISPR/Cas9 was employed to delete each of these regions in Halo-tagged PGL-1 mutants to test for P granule assembly and fertility. Immunoblots showed similar protein expression levels in PGL-1 and IFE-1 in all mutants **(Fig. S2D)**. Imaging revealed that PGL-1 ΔCtA or ΔCtB deletion mutants had normal quantity and sized PGL-1 granules compared to PGL-1 wildtype controls **(Fig. 3A,B,D)**. In contrast, the PGL-1 ΔCtC deletion mutant had fewer granules **(Fig. 3A,B)** that were significantly larger than wild type **(Fig. 3A,D)**, similar to those observed when the entire Ct region was deleted. These results suggest that PGL-1 CtC region functions in vivo to regulate P granule size and quantity. IFE-1 was also imaged to determine how the PGL-1 deletions affected protein partner recruitment. Wildtype and PGL-1ΔCtB deletion mutants formed IFE-1 puncta that colocalized with PGL-1 **(Fig. 3A,C,F)**. In contrast, PGL-1ΔCtA deletion mutants lacked IFE-1 puncta that colocalized with PGL-1 **(Fig. 3A,C,F)**, a phenotype resembling removal of the entire Ct region. While PGL-1 granule quantities were also decreased in PGL-1ΔCtC mutants **(Fig. 3A,B)**, IFE-1 puncta were still observed to colocalize with PGL-1 granules **(Fig. 3A,F).** The results support the PGL-1 CtA region being necessary for IFE-1 protein recruitment to P granules.

**Figure 3.**
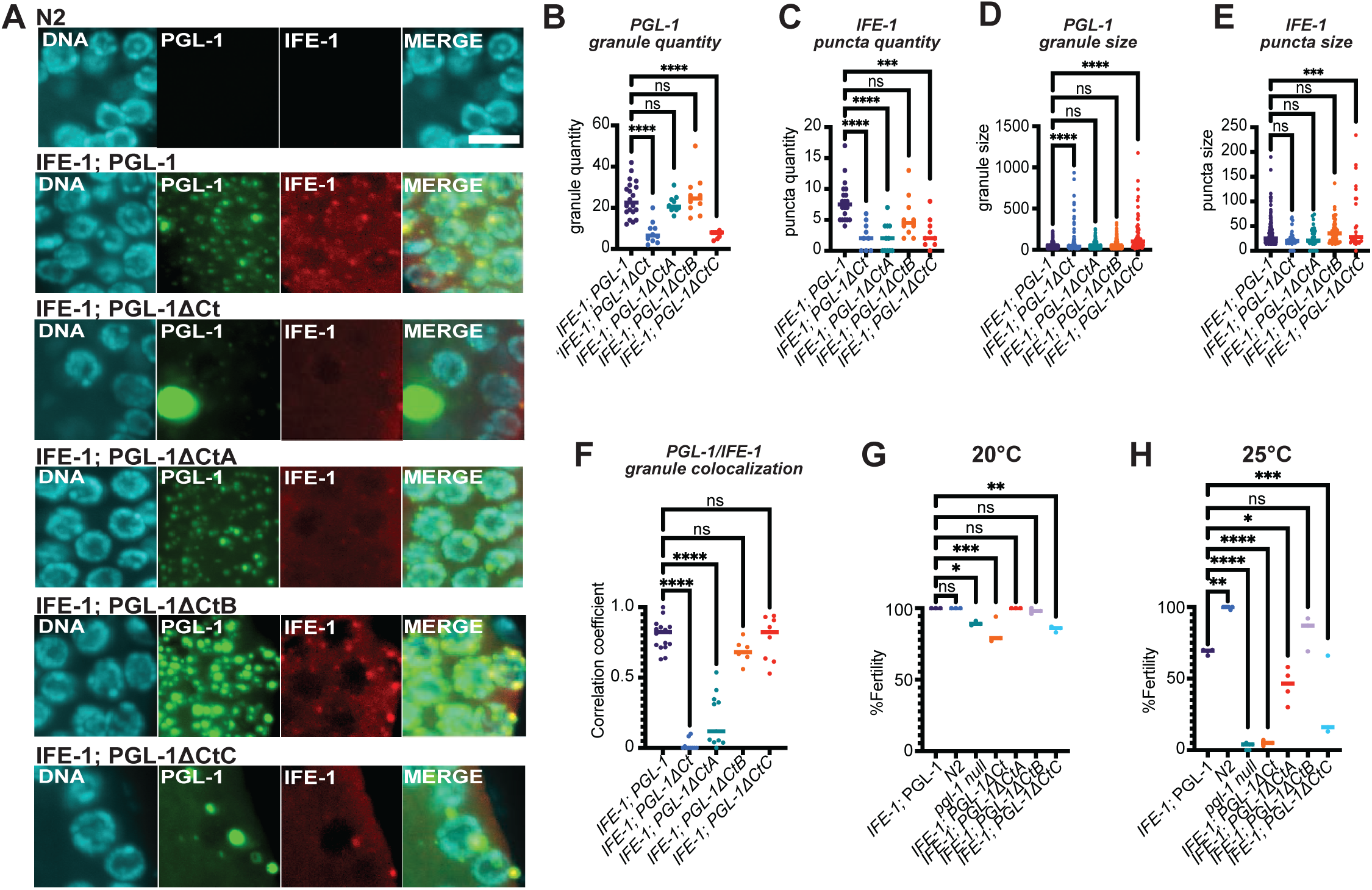
Deletions within the PGL-1 C-terminal region disrupt formation and impair fertility. (A) Confocal microscopy of adult hermaphrodite germlines. Strains expressed IFE-1::3xFLAG::mScarlet (IFE-1) with PGL-1::V5::Halo (PGL-1), PGL-1ΔCt::V5::Halo (PGL-1ΔCt), PGL-1ΔCtA::V5::Halo (PGL-1ΔCtA), PGL-1ΔCtB::V5::Halo (PGL-1ΔCtB), PGL-1ΔCtC::V5::Halo (PGL-1ΔCtC). DNA, DAPI (cyan); PGL-1, Halo Oregon Green (green); IFE-1, αFLAG (red); and a merged panel. Representative images are 10-layer z-stack maximum intensity projections. Scale bar, white, 5 μm. IFE-1; PGL-1 and IFE-1; PGL-1ΔCt images replicated from Figure 1. (**B-F**) Confocal image analyses of PGL-1 granule and IFE-1 puncta. PGL-1 granule (B) and IFE-1 puncta (C) quantification. PGL-1 granule (D) and IFE-1 (E) puncta size quantification. Correlation of IFE-1 puncta colocalizing with PGL-1 granules (F). (**G,H**) Fertility of Halo-tagged PGL-1 wildtype and mutant hermaphrodites. Worms were plated individually, propagated at 20°C (G) or 25°C (H), and scored for presence of larval progeny after 4 days. Results reported as the percentage of fertile animals relative to the total number tested in each experiment. At 20°C, fertility is significantly reduced for *pgl-1 null,* PGL-1ΔCt, and PGL-1ΔCtC compared to Halo-tagged PGL-1 animals. At 25°C, fertility was significantly impaired in all but PGL-1ΔCtB. *n =* 3 independent experiments. Ordinary one-way ANOVA statistical test was used to compare all data. ns, not significant; *, p-value <0.05. **, p-value 0.01 ***, p-value 0.001. ****, p-value <0.0001.

Fertility assays were again employed to probe the requirements of each Ct region for germ cell development. At 20°C, wild-type N2, Halo-tagged PGL-1, PGL-1ΔCtA, and PGL-1ΔCtB worms maintained high fertility while *pgl-1 (bn101) null*, PGL-1ΔCt, and PGL-1ΔCtC worms showed less fertility **(Fig. 3G).** At 25°C, PGL-1ΔCtB worms were comparable to the Halo-tagged PGL-1 control **(Fig. 3H)**, but PGL-1ΔCtA and PGL-1ΔCtC had significant fertility defects, albeit less significant than *pgl-1 null* or removal of the entire Ct region **(Fig. 3H)**. In sum, PGL-1 ΔCtA and ΔCtC mutants had defects in P granule formation and exhibited temperature-sensitive sterility. A simple explanation is that both IFE-1 recruitment to P granules and proper P granule morphology are separate functions of the PGL-1 C-terminus required for germ cell development.

IFE-1 is an eIF4E homolog that recognizes the 7-methylguanosine 5’-cap (m^7^G) of mRNAs [14]. Prior studies used m^7^G beads to capture recombinant IFE-1 and test direct PGL-1 protein binding [10]. A similar in vitro recombinant protein binding assay was used to test PGL-1/IFE-1 protein interaction (see **Methods**). As expected and shown by others [6, 10, 23], m^7^G beads enriched for recombinant IFE-1 compared to mock bead negative controls **(Fig. S3)**. Full length, recombinant PGL-1 co-eluted with enriched IFE-1 **(Fig. S3, Lane 7)**, whereas recombinant PGL-3 did not show enrichment **(Fig S3, lane 10)**, consistent with published studies [6, 10]. Prior work identified PGL-1 mutant R123E disrupted Nt domain dimerization [18], granule assembly in vitro [21]and in vivo [18], and animal fertility [18]. To test the role of PGL-1 granule assembly in IFE-1 binding, recombinant PGL-1 R123E and IFE-1 pulldowns were performed with m^7^G and mock control beads. PGL-1 R123E binding was also observed (**Fig. S3, Lane 9**), similar to wildtype protein. Thus, PGL-1 Nt dimerization or granule assembly were not required for IFE-1 interaction. Finally, recombinant PGL-1ΔCtA protein was used to test the necessity of CtA for IFE-1 binding. The removal of CtA caused a loss of PGL-1 binding **(Fig. S3, Lane 8)**. These results support that the PGL-1 CtA region is required for PGL-1/IFE-1 interaction *in vitro*.

### The PGL-1 CtA region restores granule formation and fertility when inserted into PGL-3

PGL-1 is partially redundant with PGL-3 [6], with a noted difference being that PGL-1 binds to IFE-1 while PGL-3 does not [6, 10]. Given that PGL-1 ΔCtA mutants lost IFE-1 colocalization with PGL-1 **(Fig. 3F)**, a question was whether the PGL-1 CtA peptide sequence alone was sufficient to restore function when inserted into PGL-3. To test this, the PGL-1 CtA sequence was inserted into a homologous location in an endogenous, SNAP-tagged *pgl-3* locus, generating a PGL-1/PGL-3 protein chimera **(Fig. 4A)**. Worms were crossed to express this PGL-3/PGL-1 chimera with PGL-1ΔCtA and mScarlet-tagged IFE-1 to probe rescue of IFE-1 P granule localization and germ cell function. Strikingly, addition of the PGL-1 CtA to PGL-3 rescued IFE-1 localization to P granules in a PGL-1ΔCtA background **(Fig. 4B)**. Granule quantity for PGL-1 and PGL-3 did not differ significantly across strains **(Fig. 4C,E)**, although IFE-1 puncta were reduced in the PGL-1ΔCtA mutant **(Fig. 4D)**. Puncta sizes were comparable among all mutants, with no significant differences **(Fig. 4F-H)**. Fertility assays further confirmed functional rescue. At 20°C, all strains except the *pgl-1 null* mutant maintained full fertility **(Fig. 4I)**. At 25°C, fertility dropped to ∼0% in the *pgl-1 null* strain and ∼40% in the PGL-1ΔCtA background mutant **(Fig. 4J)**. In contrast, the inclusion of the PGL-3/PGL-1 chimera restored fertility to near full-length control levels (∼70%) **(Fig. 4D)**. These results demonstrate that the PGL-1 CtA region is both necessary and sufficient to restore granule formation and fertility in *pgl-1* mutant animals.

**Figure 4.**
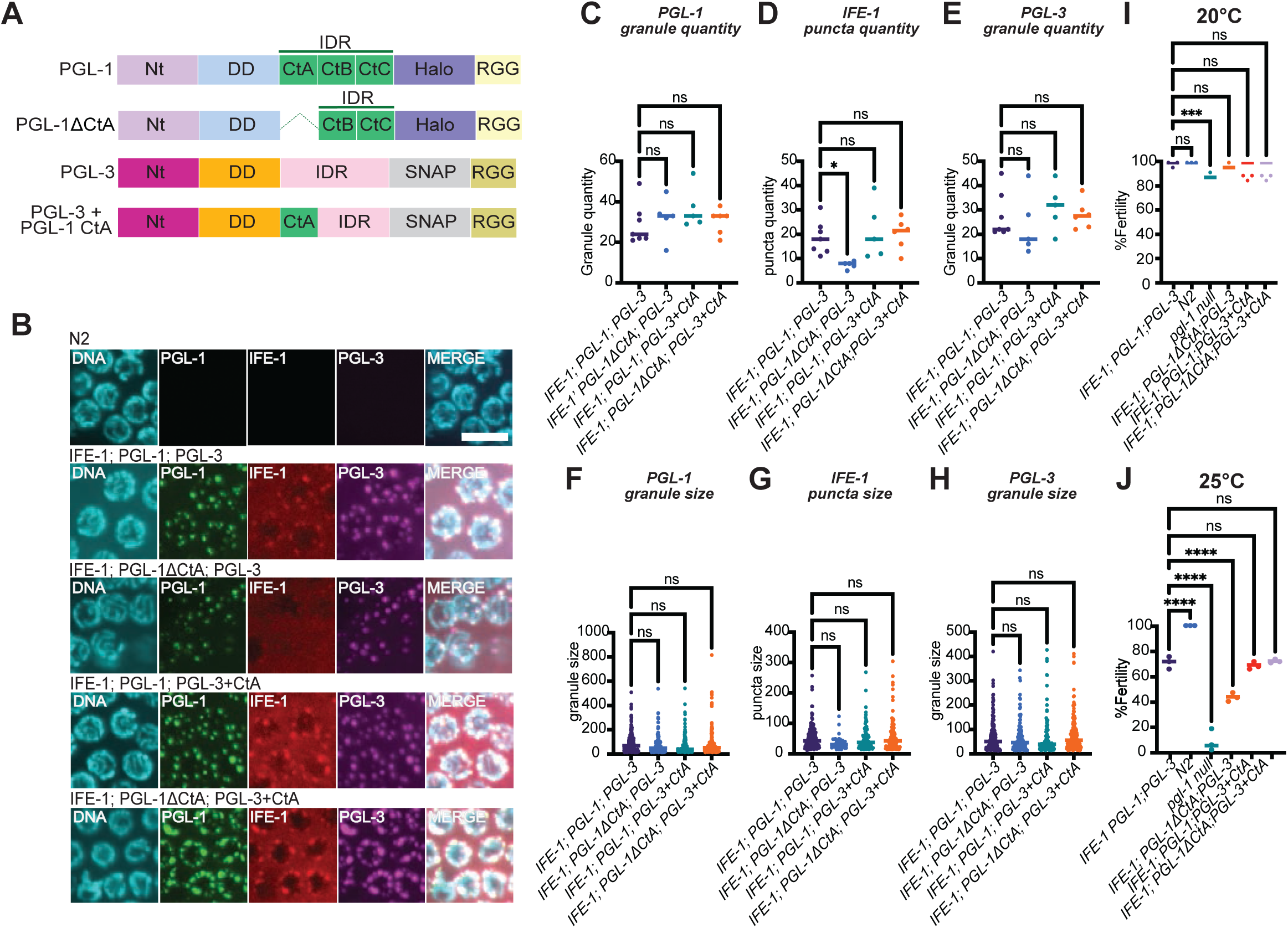
Addition of the PGL-1 CtA region to PGL-3 rescues IFE-1 localization to P granules and restores fertility. **(A)** Schematic representation of the PGL-1/PGL-3 chimera. Two dimerization domains (Nt and DD), intrinsically disordered regions (IDRs), RGG repeats, and tags are shown. The PGL-1ΔCtA is illustrated with a dashed line indicating deletion of the CtA region, while the PGL-3 + PGL-1 CtA depicts insertion of the PGL-1 CtA region (green) into the endogenous, homologous *pgl-3* region. (**B**) Confocal microscopy of adult hermaphrodite germlines. Strains expressed IFE-1::3xFLAG::mScarlet (IFE-1) with PGL-1::V5::Halo (PGL-1) or PGL-1ΔCtA::V5::Halo (PGL-1ΔCtA), and PGL-3::SNAP (PGL-3) or PGL-3::PGL-1 CtA::SNAP (PGL-3+CtA). N2 served as a wildtype control.. All strains were stained and imaged by confocal microscopy. DNA, DAPI (cyan); PGL-1, Halo Oregon Green (green); IFE-1, αFLAG (red); PGL-3, SNAP 647-SiR (magenta); and a merged panel. Representative images are 10-layer z-stack maximum intensity projections. Scale bar (white) 5 μm. (**C-E**) Quantification of PGL-1 (C) and PGL-3 (E) granule, and IFE-1 puncta (D) quantity. (**F-H**) Granule and puncta size for PGL-1 (F), PGL-3 (H), and IFE-1 (G). (**I,J**) Fertility assays of hermaphrodites carrying the indicated genotypes. Worms were plated individually at 20°C (I) or 25°C (J) and scored for larval progeny after 4 days. Results are expressed as the percentage of fertile animals relative to the total tested. At 25°C, fertility defects in PGL-1ΔCtA mutants (∼40%) were rescued by the PGL-1/PGL-3 chimera (∼70%), compared to infertility in *pgl-1* null animals (∼0%). ns, not significant; *, p < 0.05; **, p < 0.01; ***, p < 0.001.

### IFE-1 binds primarily to germline and early embryo transcripts

As an eIF4E homolog, IFE-1 should bind to the 5’ cap of mRNAs to regulate target transcripts. Prior studies sequenced IFE-1 associated transcripts (RIP-seq) [28]. To determine the targets and RNA footprint of IFE-1, enhanced cross-linking and immunoprecipitation (eCLIP, [29]) was performed by enriching for and determining the RNA fragments associated with FLAG-tagged IFE-1 **(Fig. 5A**, **Fig. S4A,B**, See **Methods)**. This data was analyzed in two ways. First, IFE-1 targets were determined by comparing peak enrichment to input samples (see **Methods)**. 883 RNAs were identified as IFE-1 targets in this manner. tRNA was the most prevalent RNA identified (∼83%), whereas protein coding mRNA was less abundant (4%) **(Fig. S4C)**. Most targets were located on the X chromosome (∼40%) **(Fig. S4D)**, a surprising result given that the X chromosome is silenced in most of the adult germline [30–32]. Comparison of IFE-1 eCLIP targets to published RIP-seq results [28] revealed little overlap **(Fig. S4E)**.

**Figure 5.**
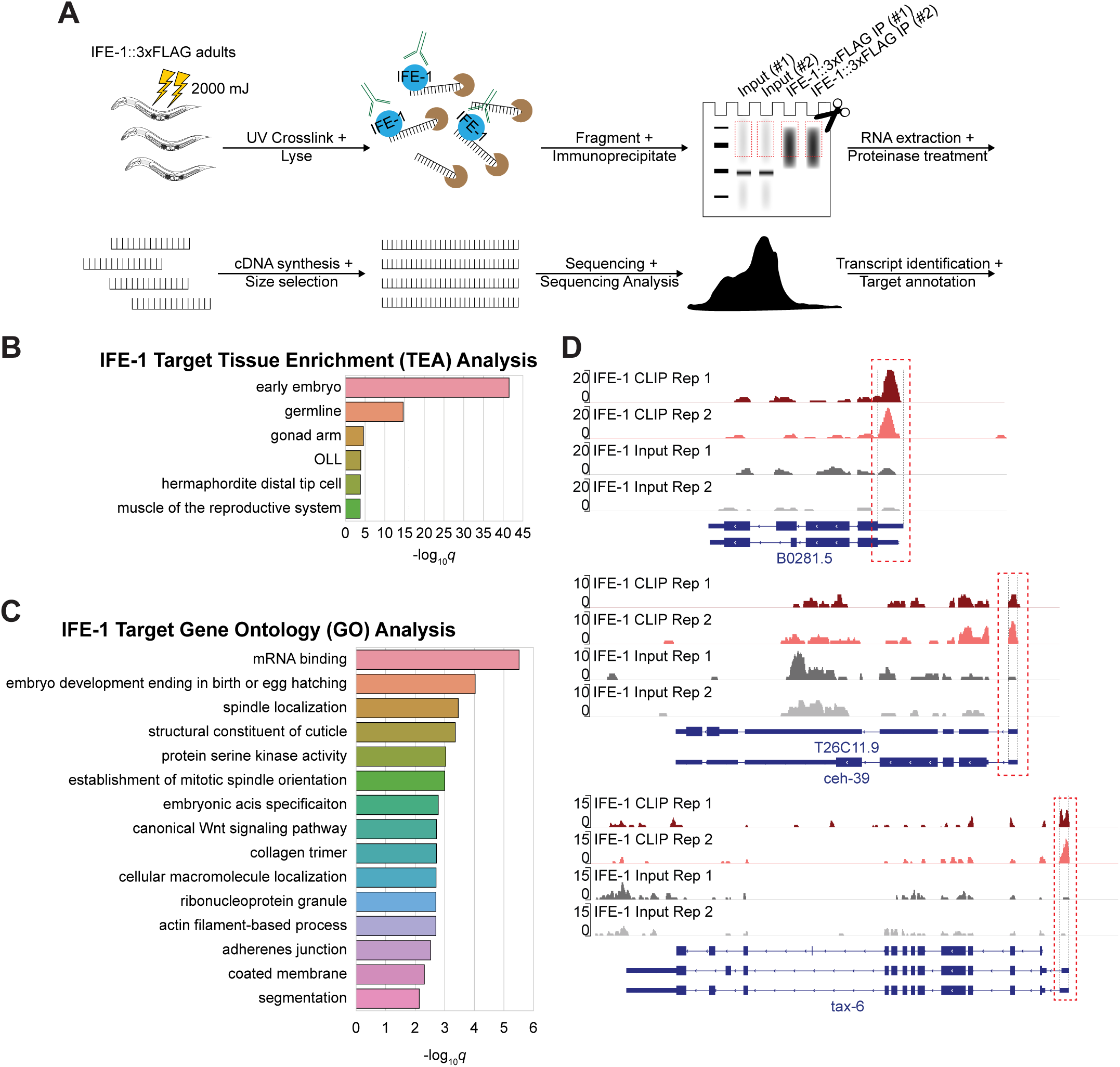
IFE-1 binds transcripts involved in mRNA regulation and fertility. **(A)** IFE-1 eCLIP workflow. Synchronized young adult hermaphrodite worms expressing IFE-1::3xFLAG::mScarlet (IFE-1::3xFLAG) were UV crosslinked, lysed, and IFE-1 immunoprecipitated with a FLAG antibody. Bound RNA was separated by SDS-PAGE, processed, and sequenced to identify IFE-1 targets and footprints. (**B**) Tissue enrichment analysis (TEA) performed on top 25% of IFE-1 ranked mRNA targets identified in 5’ ratio analysis. The early embryo and germline associated genes are highly enriched. (**C**) Gene ontology analysis (GO) performed on top 25% of IFE-1 ranked mRNA targets identified in 5’ ratio analysis. (**D**) Genome browser views of highly ranked mRNA transcript examples identified in 5’ ratio analysis. Transcripts *B0281.5*, *ceh-39*, and *tax-6* shown. IP replicates in red, input replicates in grey. Red dotted box denotes 5’ untranslated region (UTR) with associated peaks.

The second eCLIP analysis strategy leveraged the functional knowledge that IFE-1 should specifically interact with the 5’ RNA cap. Therefore, true mRNA targets should have footprints at their 5’ ends. A ratio statistic was computed for every mRNA transcript by comparing enrichment in 20% of the 5’ end to the remainer of the transcript (see **Methods)**. These ratios were ranked and filtered to remove small RNAs and other background. With this strategy, 982 transcripts represented the top 25% and used in downstream analyses **(Supplementary Table 1)**. The targets represented genes from all chromosomes **(Fig. S4F)** and little overlap was observed when compared to previous studies [28] **(Fig. S4G)**. This limited overlap may be due to methodology differences. As P granules contain dense concentrations of RNA, the use of sequencing methods without crosslinking or 5’ end enrichment may lead to identification of non-specific interactions. Regardless, highly ranked transcripts in this study’s eCLIP analysis were those found in the *C. elegans* early embryo and germline **(Fig. 5B)**. Gene ontology analysis enriched for processes involved in RNA regulation and reproduction **(Fig. 5C)**. Collectively, this analysis identifies IFE-1 target transcripts in the adult germline.

### Loss of IFE-1 recruitment to PGL-1 leads to derepression of IFE-1 mRNA targets under temperature stress

In the adult germline, IFE-1 is postulated to activate its mRNA target transcripts [28], but P granules are proposed to be sites of mRNA repression [9, 17, 18]. To directly clarify this discrepancy, top gene targets with 5’ UTR peaks **(Fig. 5D)** were modified by CRISPR/Cas9 to add a SNAP tag to their protein coding regions, generating protein expression reporters to genes *B0281.5, tax-6*, and *ceh-39. B0281.5* encodes a homolog to the human potassium channel tetramerization domain containing 7 (KCTD7) with no known tissue specificity [33, 34]. *tax-6* encodes a calcineurin A subunit with expression detected in tail neurons and rays in male *C. elegans* [35]. *ceh-39* is a homeobox gene is located on chromosome X, involved in neuronal development, and can be found in the nuclei of young embryos and in the later proximal germline [36]. The reporter worms were crossed with *pgl-1 null*, and Halo-tagged PGL-1 wildtype and mutant worms to probe the connection between reporter expression and IFE-1/PGL-1 assembly.

Worms were propagated at 20°C or 25°C, the temperature associated with the *pgl-1 null* phenotype, followed by imaging to detect germline protein expression. SNAP-tagged proteins have been successfully used to visualize proteins in the *C. elegans* germline [18] with low background in wild-type N2 negative controls **(Fig. 6A)**. PGL-1 appeared as granules at the nuclear periphery in all Halo-tagged PGL-1 strains **(Fig. 6A,C)**, consistent with prior observations **(Fig. 1**, **Fig. 3)**. In CEH-39 reporter worms, no expression of SNAP-tagged CEH-39 could be observed with the reporter alone or with Halo-tagged PGL-1 at 20°C **(Fig. 6A,B)**, while modest CEH-39 reporter expression could be detected in *pgl-1 null* worms at both 20°C and 25°C **(Fig. 6A-D)**. Reporter worms with PGL-1ΔCtA lacked CEH-39 expression at 20°C but had detectable expression at 25°C **(Fig. 6A,C)**. In B0281.5 reporter worms, no expression of the reporter was detectable at 20°C in any of the genetic backgrounds tested **(Fig. S5A,B)**. B0281.5 protein could be observed at 25°C in both *pgl-1 null* and PGL-1ΔCtA worms **(Fig. S5C,D)**, albeit only quantifiably significant in the *pgl-1 null* worms **(Fig. S5D)**. The low level of protein expression most likely made reporter quantitation challenging over background. SNAP-tagged TAX-6 protein could not be observed in the germline in any of the genetic backgrounds and temperatures tested **(Fig. S6A-D)**, despite detectable protein expression in whole worm immunoblots **(Fig. S6E)**. The lack of germline expression compared to the other reporters is unclear. *tax-6* mRNA may be under additional levels of post-transcriptional gene regulation, such as by 3’UTR binding proteins [37] or small RNA regulation [38, 39]. It also may be challenging to quantify as a cytoplasmic protein, or it may be a false positive IFE-1 mRNA target. Regardless, the lack of detectable TAX-6 germline protein expression in *pgl-1 null* worms provides evidence that the observed protein expression of the CEH-39 and B0281.5 reporters is not merely an aberrant artifact due to sterile germlines. In sum, these findings support the model that IFE-1 represses its target transcripts. Some IFE-1 target genes were derepressed with the loss of PGL-1 binding, suggesting that IFE-1 recruitment to P granules is necessary for its repressive function in vivo (see **Discussion**).

**Figure 6.**
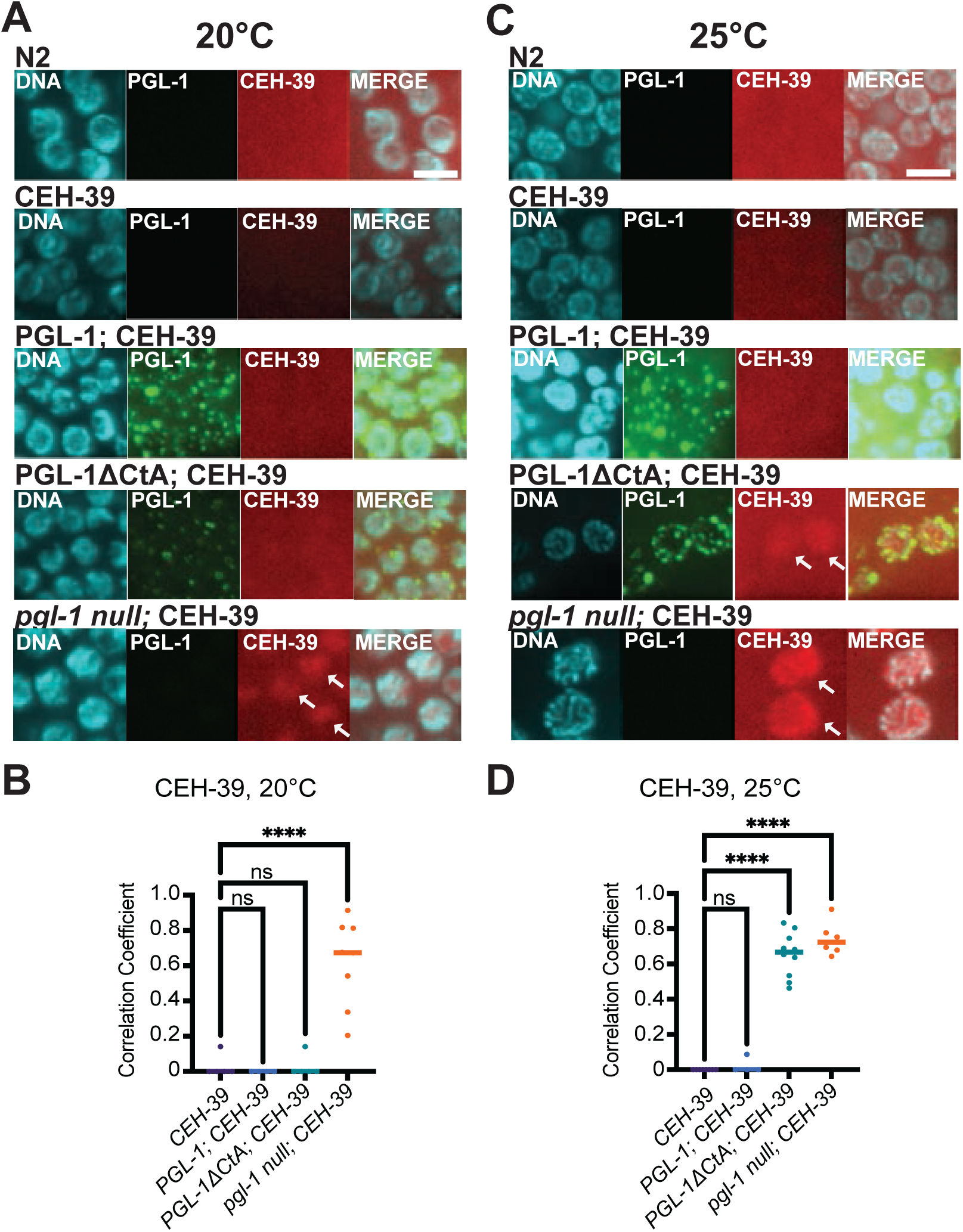
Germline CEH-39 protein expression increases when IFE-1 assembly into P granules is disrupted. Confocal microscopy images of adult hermaphrodite germlines. Strains expressed CEH-39::SNAP (CEH-39) with PGL-1::V5::Halo (PGL-1), PGL-1ΔCtA::V5::Halo (PGL-1ΔCtA), or *pgl-1*(bn101) (*pgl-1 null)*. Adult germlines were imaged by confocal microscopy. DNA, DAPI, cyan; PGL-1, Halo Oregon Green, green; CEH-39, SNAP TMR, red; and a merged panel. White scale bar, 5 μm. All images and analyses performed on 10-layer z-stack maximum intensity projections. (**A**) Representative images of CEH-39 worms in PGL-1, PGL-1ΔCtA, and *pgl-1 null* genetic backgrounds, propagated at 20°C. CEH-39 expression is observed only in the *pgl-1 null (bn101)* background, indicated by white arrows. (**B**) Quantification of CEH-39 signal colocalization with DNA at 20°C. (**C**) Representative images of the SNAP-tagged CEH-39 worms previously described, propagated at 25°C. White arrows highlight CEH-39 expression with PGL-1ΔCtA or *pgl-1 null* genetic backgrounds. (**D**) Quantification of CEH-39 colocalization with DNA at 25°C. ns, not significant; ****, p-value <0.0001.

## Discussion

This study explored how the PGL-1 P granule assembly component interacts with the IFE-1, an eIF4E cap-binding protein to promote *C. elegans* germ cell development. Using microscopy, biochemistry, and molecular genetic methods, the investigation made three key findings. First, it determined a previously uncharacterized role for the PGL-1 Ct region in IFE-1 recruitment and granule formation. Second, it determined direct RNA targets bound to IFE-1. Third, it provided direct evidence that IFE-1 associated mRNAs are repressed and that this repressive function is dependent on PGL-1 association. Together, these findings support a model where IFE-1 is recruited to P granules by directly binding the PGL-1 C-terminus to repress its mRNA targets for germ cell development **(Fig. 7)**.

**Figure 7.**
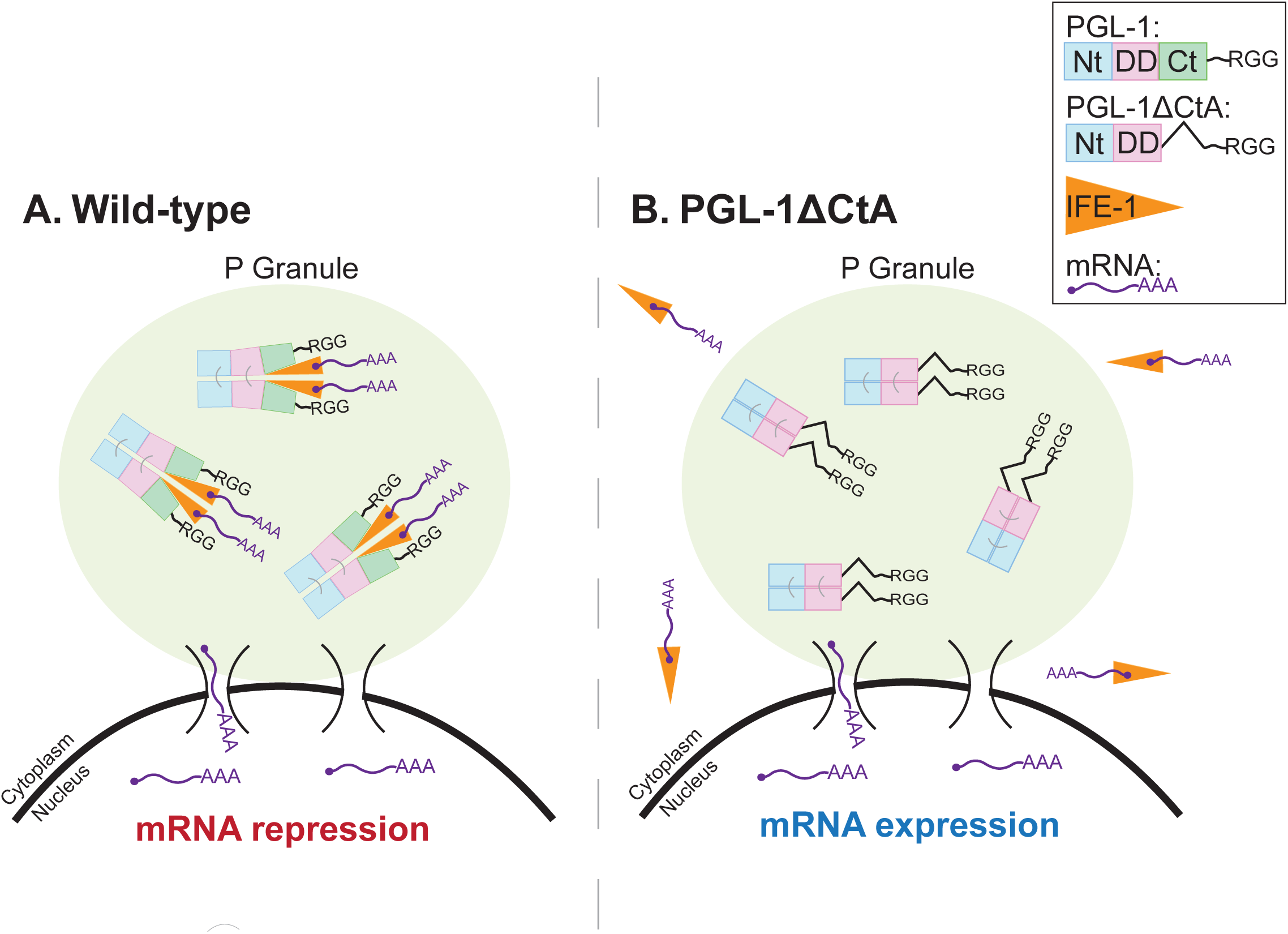
Model of IFE-1 assembly into P granules for mRNA target repression. **(A)** P granules assemble at the nuclear pore with PGL-1 and IFE-1 protein. For simplicity, PGL-1 is shown self-dimerizing via its Nt (blue) and DD (pink) domains. IFE-1 (orange) localizes to P granules via direct binding to PGL-1 CtA (green) to repress its target mRNAs. (**B**) Removal of PGL-1 CtA prevents proper IFE-1 assembly into P granules, permitting target mRNA expression.

### IFE-1 functions as a negative regulator in protein expression

eIF4E recognizes the m^7^G cap on the 5’ ends of mRNAs and plays a central role in regulating messenger ribonucleoproteins (mRNPs) localization, storage, turnover, translation, and mRNA repression [14]. As an activator, eIF4E assembles with the eIF4G scaffold protein to coordinate translation initiation of target transcripts. In metazoans, this eIF4E/eIF4G interaction is conserved and negatively regulated by 4E-binding proteins (4E-BPs), which can compete with eIF4G for a similar binding site of eIF4E [14]. In germ cells and embryos, specific 4E-BPs have been shown to block eIF4G binding in a similar manner [40–43]. This mode of translational repression by 4E-BPs is conserved across species, including *Drosophila* and mammals [44].

PGL-1 was previously identified as 4E-BP that interacts with IFE-1 [7]. Other studies have proposed that IFE-1 is a translational activator based on an increase in target gene expression in *ife-1* null animals [28]. While IFE-1 binds to m^7^G-capped mRNAs, consistent with a canonical role in promoting translation, the findings reported here suggest that IFE-1 assembly with PGL-1 causes target transcript repression, at least within developing adult germ cells. These two models are not mutually exclusive, as removal of the eIF4E bound to these transcripts would expose the mRNA 5’ end for turnover [45]. As observed in other systems [44], the 4E-BPs bound to IFE-1 may dictate its molecular function. How PGL-1 represses IFE-1 mRNA targets is not fully understood. 4E-BPs are classically translational repressors [14], but prior work noted that localizing transcripts to P granules causes mRNA turnover [18]. Alphafold3 [46] modeling predicts that PGL-1 CtA directly binds to IFE-1 opposite its m^7^G binding site **(Fig. S7)**, similar to other 4E-BPs [47, 48]. PGL-1 may compete with eIF4G binding to balance translational repression and activation, respectively. Additional structural and biochemical studies will be necessary to clarify how IFE-1 assembles in P granules and represses its target transcripts.

IFE-1 is expressed throughout all stages of germline development and required for spermatogenesis [10–13]. It directly associates with PGL-1 until the clearance of P granules in primary spermatocytes through spermatid budding [10]. The loss of P granules coincides with the transcriptional activation and translation of sperm-related transcripts [49]. P granules also play a key role in safeguarding germline identity by preventing the translation of transcripts associated with somatic cell fates [17, 50] and limiting the presence of sperm-specific mRNAs in adult hermaphrodites [49]. This work identified that IFE-1 primarily associates with early embryo genes in adult germline, and that IFE-1 localization to P granules correlated with gene target repression. A spectulation is that these genes should also be repressed in developing adult germ cells to permit proper gamete formation. Of note, *ife-1 null* mutants are sterile [11, 12] but PGL-1 ΔCtA mutants only partly sterile. IFE-1-associated transcripts may be regulated independently of P granules through secondary repressive mechanisms. Further work investigating the regulation of IFE-1 target transcripts and the role of IFE-1 in spermatogenic animals will clarify how P granules function in germ cell development.

### P granules as sites for mRNA repression

Germ granules are specialized RNA-protein condensates in germ cells that regulate RNA fate, including storage, translation, and processing [4]. P granules localize on the cytoplasmic face of nuclear pore complexes [5]. The proximity likely enables germ granules to monitor nascent transcripts exiting the nucleus, as evidenced by the accumulation of these transcripts in perinuclear P granules [9]. Transcriptome profiling of germlines lacking P granule assembly components revealed widespread misregulation, including inappropriate expression of transcripts typically restricted to sperm or somatic lineages [49, 50]. These findings suggest that P granules play a crucial role in germ cell differentiation by repressing ectopic gene expression. How do P granules mediate such repression? Prior work showed that artificial tethering PGL-1 to reporter mRNA repressed its expression, and that this repression was dependent on PGL-1 Nt dimerization [18], a mutation that also affected P granule assembly [18]. While the study provided evidence for granule assembly being required for mRNA repression, concerns for the work included the artificiality of mRNA tethering, the sterility observed with PGL-1 tethering, and that molecular attributes other than granule assembly may be affected in protein dimer disruption. This current work determined that endogenous tethering of PGL-1 to IFE-1 represses its associated transcripts, and it identified a PGL-1 mutant that affects IFE-1 assembly and its associated repression while maintain PGL granule formation. Thus, two lines of evidence now support P granules as sites of mRNA repression and the requirement of proper P granule assembly for this repression. The PGL-1 C-terminal tail is responsible for recruiting IFE-1. As dozens of mRNA-associated proteins have been identified in P granules [5], other binding proteins may be recruited by the PGL C-termini for mRNA repression. Removal of a separate PGL-1 C-terminal region caused aberrant granule formation and was at least partially attributable to PGL-1’s biological function. Regions in the C-terminus may associate with other factors that promote proper condensate formation. Future work should identify other binding partners that facilitate P granule’s assembly and repressive functions.

### P granules and RNA-protein condensate form and function

Thus far, many RNA-protein and germ granules have been observed to have phase separation properties. For example, Balbiani bodies are present in many animal species, including mammals [51, 52]. They form adjacent to the nucleus of primary oocytes and are composed of organelles, specific proteins, and RNAs. *Drosophila melanogaster* germ granules are electron dense, RNA-protein condensates that function as key hubs of RNA regulation and are essential for germline specification [53, 54]. Stress granules form in the cytoplasm in response to environmental stress, sequestering untranslated mRNAs [55] until they dissolve once the cellular stress is alleviated [56]. In these cases, condensates form stable, solid-like states that physically trap mRNAs, unsurprisingly repressing and storing them for later use. P granules were the first RNA-protein condensates reported to behave like liquid droplets [6, 10, 11]. The function of liquid condensates is not intuitive, as an mRNA in liquid may diffuse out for protein expression. This work supports P granules as active biochemical hubs of mRNA regulation, where mRNA-binding proteins that localize to P granules leads to repression of its targets. P granules are also well known germ granule hubs that interact with Z granules, SIMR foci, mutator foci, and other condensates for small RNA processing and post-transcriptional regulation [5]. As the biochemistry of P granules unfolds, these other granule players will likely function in collaborative roles with this liquid condensate to promote germ cell development.

## Acknowledgements

The authors thank members of the Aoki and Hundley lab for thoughtful discussions. We also thank Douglas Rusch from Indiana University Bloomington for bioinformatic analysis. B.B. was supported by the Indiana Medical Scientist Training Program (MSTP) summer research program. B.Y. and H.H. are supported by R35GM156459. S.T.A., C.H.S., C.B., and this work was funded by Indiana University’s start-up funds and its Precision Health Initiative, the Showalter Trust, and the NIH/NIGMS (R35GM142691).

## Methods

### Nematode Strains and Maintenance

Nematodes were cultured on NGM plates (25 mM KPO_4_ pH 6.0, 5 mM NaCl, 1 mM CaCl_2_, 1 mM MgSO_4_, 2.5 mg/ml tryptone, 5 µg/ml cholesterol, 1.7% agar) with HB101 bacteria as food source as described previously [57]. All strains were grown from 15°C to 20°C. To synchronize populations at the first larval stage, worms were bleached and allowed to hatch overnight in M9 buffer using standard protocols [58]. In temperature experiments, worms were exposed to specific temperatures as L4 larvae and allowed to propagate. F1 worms were collected, stained, and imaged as described below.

The following strains were used in this study:

| Strain Name | Description | Genotype |
| --- | --- | --- |
| N2 | wildtype |  |
| SS579 | *pgl-1 null* | *pgl-1(bn101) IV* |
| CDE78 | *Halo-tagged PGL-1* | *pgl-1 (ddc26 [pgl-1::V5::Halo]) IV* |
| CDE79 | *mScarlet-tagged IFE-1* | *ife-1 (ddc27 [ife-1::3xFLAG::mScarlet (introns)]) III* |
| CDE102 | *Halo-tagged PGL-1ΔCtA* | *pgl-1 (ddc44 [pgl-1 del aa. 452-553::V5::Halo]) IV* |
| CDE103 | *Halo-tagged PGL-1ΔNt* | *pgl-1 (ddc45 [pgl-1 del aa. 10-213::V5::Halo]) IV* |
| CDE104 | *Halo-tagged PGL-1ΔCt* | *pgl-1 (ddc46 [pgl-1(del aa. 452-672)::V5::Halo]) IV* |
| CDE106 | *Halo-tagged PGL-1ΔCtC* | *pgl-1 (ddc48 [pgl-1(del aa. 611-672)::V5::Halo]) IV* |
| CDE108 | *Halo-tagged PGL-1ΔDD* | *pgl-1 (ddc50 [pgl-1(del aa. 214-451)::V5::Halo]) IV* |
| CDE100 | *Halo-tagged PGL-1RGG* | *pgl-1 (ddc42 [pgl-1::V5::HaloOPT del. 673-730]) IV* |
| CDE129 | *Halo-tagged PGL-1ΔCtB* | *pgl-1 (ddc67 [pgl-1(del aa. 554-610)::V5::Halo]) IV* |
| CDE130 | *mScarlet-tagged IFE-1; Halo-tagged PGL-1ΔCtA* | *ife-1 (ddc27 [ife-1::3xFLAG::mScarlet (introns)]) III; pgl-1 (ddc44 [pgl-1 del aa. 452-553::V5::HaloOPT]) IV* |
| CDE131 | *mScarlet-tagged IFE-1; Halo-tagged PGL-1* | *ife-1 (ddc27 [ife-1::3xFLAG::mScarlet (introns)]) III; pgl-1 (ddc26 [pgl-1::V5::Halo]) IV.* |
| CDE132 | *mScarlet-tagged IFE-1; Halo-tagged PGL-1ΔRGG* | *ife-1 (ddc27 [ife-1::3xFLAG::mScarlet (introns)]) III; pgl-1 (ddc42 [pgl-1::V5::Halo del aa. 673-730]) IV* |
| CDE133 | *mScarlet-tagged IFE-1; Halo-tagged PGL-1ΔCtB* | *ife-1 (ddc27 [ife-1::3xFLAG::mScarlet (introns)]) III; pgl-1 (ddc67 [pgl-1 del aa. 554-610)::V5::Halo]) IV* |
| CDE134 | *mScarlet-tagged IFE-1; Halo-tagged PGL-1ΔNt* | *ife-1 (ddc27 [ife-1::3xFLAG::mScarlet (introns)]) III; pgl-1 (ddc45 [pgl-1 del aa. 10-213::V5::Halo]) IV* |
| CDE135 | *mScarlet-tagged IFE-1; Halo-tagged PGL-1ΔDD* | *ife-1 (ddc27 [ife-1::3xFLAG::mScarlet (introns)]) III; pgl-1 (ddc50 [pgl-1(del aa. 214-451)::V5::Halo]) IV* |
| CDE97 | *M01E5.4 null* | *M01E5.4 (ddc39 [M01E5.4::mScarlet with 62 bp deletion]) I* |
| CDE98 | *mScarlet-tagged M01E5.4* | *M01E5.4 (ddc40 [M01E5.4::mScarlet]) I* |
| CDE150 | *SNAP-tagged PGL-3* | *pgl-3 (ddc74 [pgl-3::SNAP]) IV* |
| CDE169 | *SNAP-tagged PGL-3 with PGL-1 CtA* | *pgl-3 (ddc79 [pgl-3 + PGL-1 CtA::SNAP] V* |
| CDE153 | *SNAP-tagged B0281.5* | *B0281.5 (ddc75 [B0281.5::SNAP::OLLAS] II* |
| CDE154 | *SNAP-tagged B0281.5; Halo-tagged PGL-1* | *B0281.5 (ddc75 [B0281.5::SNAP::OLLAS] II.; pgl-1 (ddc26 [pgl-1::V5::Halo]) IV.* |
| CDE155 | *SNAP-tagged B0281.5; Halo-tagged PGL-1ΔCtA* | *B0281.5 (ddc75 [B0281.5::SNAP::OLLAS] II; pgl-1 (ddc44 [pgl-1 del 452-553::V5::HaloOPT]) IV* |
| CDE163 | *SNAP-tagged B0281.5; pgl-1 null* | *B0281.5 (ddc75 [B0281.5::SNAP::OLLAS] II; pgl-1(bn101) IV.* |
| CDE156 | *SNAP-tagged TAX-6* | *tax-6 (ddc78 [tax-6::SNAP OLLAS]) IV* |
| CDE157 | *SNAP-tagged TAX-6 Halo-tagged PGL-1* | *tax-6 (ddc78 [tax-6::SNAP OLLAS]) IV; pgl-1 (ddc26 [pgl-1::V5::Halo]) IV.* |
| CDE170 | *SNAP-tagged TAX-6 pgl-1 null* | *tax-6 (ddc78 [tax-6::SNAP OLLAS]) IV; pgl-1(bn101) IV* |
| CDE158 | *SNAP-tagged TAX-6 Halo-tagged PGL-1ΔCtA* |  |
| CDE164 | *SNAP-tagged CEH-39* | *ceh-39 (ddc76 [ceh-39::SNAP::OLLAS] X* |
| CDE165 | *SNAP-tagged CEH-39; Halo-tagged PGL-1* | *pgl-1 (ddc26 [pgl-1::V5::Halo]) IV; ceh-39 (ddc76 [ceh-39::SNAP::OLLAS] X* |
| CDE167 | *Halo-tagged PGL-1ΔCtA; SNAP-tagged CEH-39* | *pgl-1 (ddc44 [pgl-1 del aa. 452-553::V5::Halo]) IV, ceh-39 (ddc76 [ceh-39::SNAP::OLLAS] X* |
| CDE168 | *PGL-1 null; SNAP-tagged CEH-39* | *pgl-1(bn101) IV; ceh-39 (ddc76 [ceh-39::SNAP::OLLAS] X* |

### Microscopy

Animal gonad dissection and imaging were performed as previously described [59]. Briefly, adult worms were collected into petri dishes containing M9 + 0.1% Tween-20 and 0.25 mM Levamisole. Gonads were extruded and collected into Eppendorf tubes with PBS-T (PBS + 0.1% Tween-20), washed three times to remove excess bacteria, fixed for 10 minutes with 1% paraformaldehyde + 0.1% Triton-X at room temperature, and incubated with combinations of anti-FLAG (M2 (mouse) (1:2000) Millipore-Sigma; Burlington, MA) for IFE-1 visualization, 100 nM Halo Oregon Green (Promega) for PGL-1 visualization, and 1 µM SNAP 647-SiR (Promega) for SNAP visualization, then placed overnight in cold room with rotation. After the incubation period, samples were washed three more times with PBS-T and incubated with combinations of fluorophore-labeled secondary Alexa 555 donkey anti-mouse antibody (1:2000) (Invitrogen, Carlsbad, CA) and DAPI 0.5 µg/ml (1:2000) (Invitrogen, Carlsbad, CA) for one hour at room temperature with rotation. Stained gonads were mounted onto a slide with Vectashield (Vector Laboratories, Burlingame, CA) and imaged on a Zeiss AxioObserverZ1 modified by 3i (www.intelligent-imaging.com) for confocal microscopy with an image stack ranging from 60-100 slices. All images were analyzed by ImageJ/Fiji Bio-Formats plugins (National Institutes of Health) [60]. Within any set of comparable images, the image capture and scaling conditions are identical. The cyan/blue channel included a 405-emission filter, a green channel included a 488**-**emission filter, a red 560-emission filter, and red 647-emission filter. For each CRISPR modified animal, the distal gonad regions were imaged and analyzed. For each condition, a minimum of two biological replicates were performed.

### Image analysis

Extruded germlines were imaged using a Zeiss AxioObserverZ1 modified by 3i (www.intelligent-imaging.com) confocal microscope and saved as .sld files. Z-stack imaging was performed, and a 10-layer stack was generated using Fiji (ImageJ) to enhance granule visualization and quantification. The z-projection of the stacked file was set to “max intensity,” and the brightness and contrast of each channel was adjusted with designated maximum and minimum intensity levels to optimize contrast and ensure accurate granule detection. DAPI was used as a nuclear marker (blue channel), Halo-Oregon Green labeled PGL-1 (green channel), and mScarlet tagged IFE-1 (red channel). Individual channels were saved separately as PNG files, along with a merged image combining all three channels to facilitate the analysis of granule localization. Scale bars were added to figures containing merged channels. Granule and puncta quantification was performed by setting intensity thresholds to 3% for PGL-1 and 1.5% for IFE-1, respectively. A single round of image smoothing was applied, and granules were quantified within a defined rectangular region of interest (ROI) (coordinates: 252, 1004, width: 154, height: 156). Particles smaller than 14 pixels were excluded from the analysis.

### Reporter Analysis

Ten-layer Z-stack images were imported into Fiji (ImageJ), and the JACoP plugin was used to quantify colocalization. A region of interest (ROI) was defined using the coordinates (X: 252, Y: 1004; width: 154, height: 156). SNAP-tagged reporters were overlaid onto the DNA channel, and thresholds were manually adjusted to exclude background signal. Manders’ correlation coefficient was used to assess colocalization, as it focuses on co-occurrence by measuring the proportion of signal in one channel that overlaps with the signal in the other. This approach is particularly well-suited for evaluating colocalization because it does not rely on signal intensity correlation but instead captures spatial overlap between markers.

### CRISPR/Cas9 Gene Editing

A co-conversion CRISPR technique was implemented in our model following a protocol generated by the Fire lab [61]. Briefly, worms were injected with a target CRISPR-Cas-9 RNA (crRNA) or plasmid expressing a Cas9-scaffold with tandem target sequence RNA (sgRNA) to a gene of interest, a target crRNA to *unc-58*, a scaffolding tracrRNA, recombinant Cas9 protein, and a *unc-58* repair DNA oligo that inserted a dominant mutation [62]. For large edits, such as fluorescent protein tag sequences, we generated double-stranded DNA repair templates by amplifying Halo by polymerase chain reaction using specific oligos containing homology arms of approximately 50 base pairs (bp). For single-nucleotide modifications or deletions, we used single-stranded oligonucleotides containing homology arms of approximately 35 bp as repair templates. F1s displaying the co-injection marker phenotype underwent additionally screened by a combination of PCR without or with restriction enzyme digest to identify those with the repair of interest. F2s were PCR screened to identify homozygous alleles, and the PCR product sequenced to confirm proper repair. The transgenic strain was crossed to the N2 strain for two generations to remove potential off target mutations. All guide RNAs and oligos were obtained commercially. The primer sequences for crRNA and amplifying the repair template are listed below:

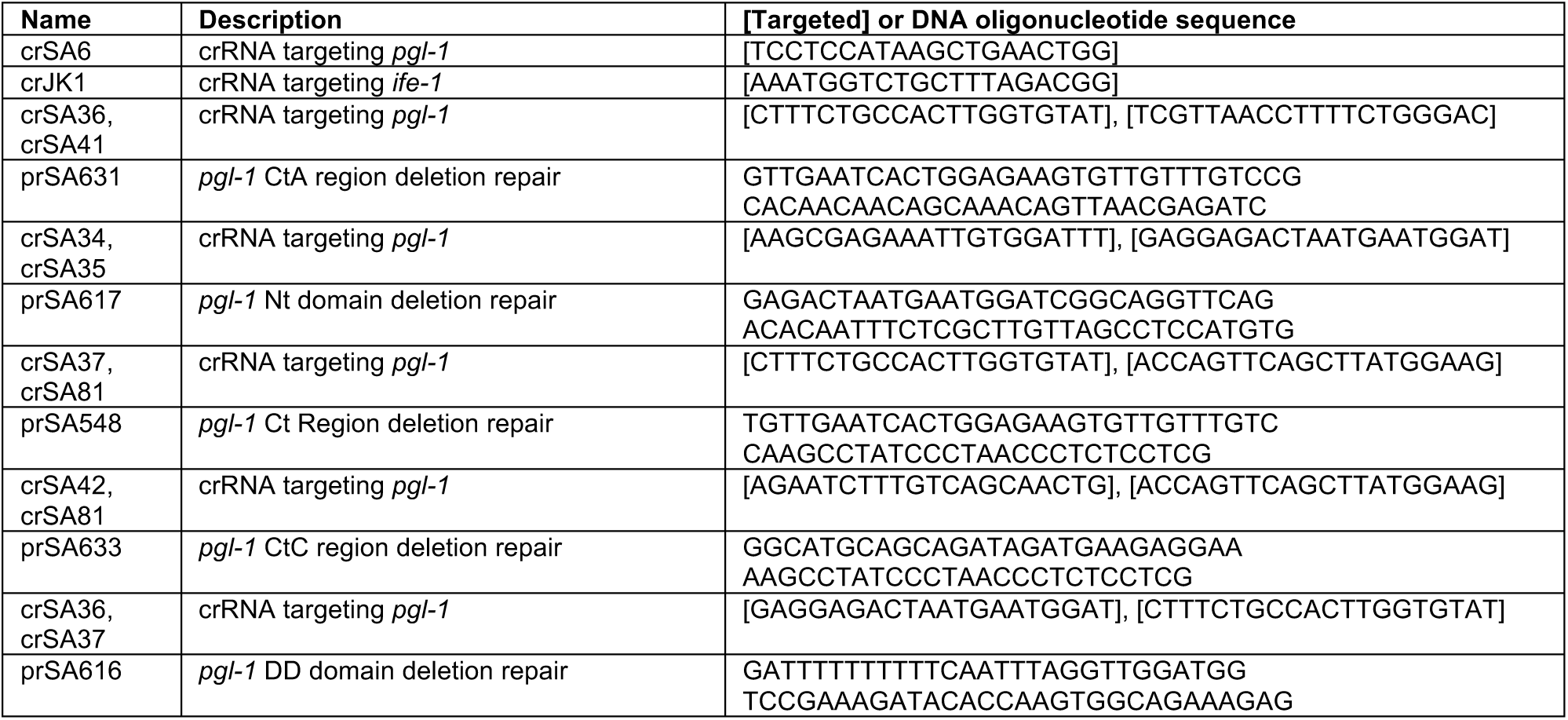
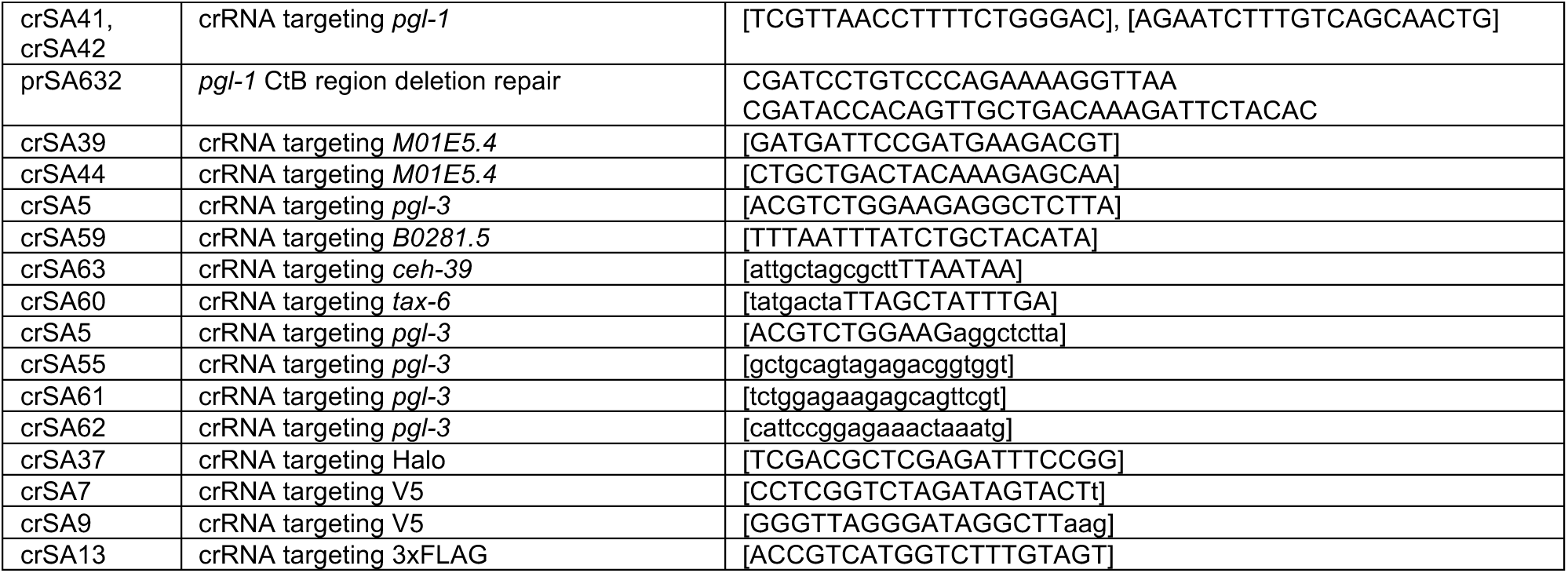

### C. elegans immunoblots

Synchronized populations of *C. elegans* adult hermaphrodites were harvested, washed three times in PBS-T, and stored in -20°C with 5x SDS buffer (250 mM Tris pH 6.8, 25 mM EDTA pH 8.0, 25% glycerol, 5% SDS, 500 mM beta-mercaptoethanol). The samples were then thawed, boiled for 10 minutes at 90°C and loaded onto SDS-PAGE gels. Proteins were transferred to a PVDF membrane (BioRad, 227 #1620264) using the BioRad TransBlot Turbo System (BioRad, #1704150). Membranes were blocked for 1 hour at room temperature with PBS-T buffer containing 5% non-fat milk, incubated overnight with primary antibodies on rotation at 4°C, and incubated for 2 hours at room temperature with HRP-conjugated antibodies. The following antibodies or protein-HRP conjugates were used for this study: V5 (E9H8O) Mouse mAb #80076 (1:1000) (Cell Signaling), GAPDH (D4C6R) Mouse mAb #97166 (1:5000) (Cell Signaling), SNAP/CLIP monoclonal (6F9) Rat Ab (1:1000)(ChromoTek), H3 (#9715) Rabbit Ab (1:1000)(Cell Signaling), β-Actin (8H10D10) Mouse mAb #3700 (1:1000)(Cell Signaling).

### Fertility assay

Fertility assays were conducted in incubators set to 20°C and 25°C. Populations were exposed to these temperatures from the L4 stage and allowed to propagate for one generation. F1 progeny were plated onto 24-well plates containing *E. coli* strain HB101 and returned to the respective incubators for four days. Fertility was assessed by scoring wells for larval growth, with only wells containing live progeny counted as fertile. All fertility experiments were repeated at least 2-3 times and reported as percent fertile (% fertile = n fertile/n total).

### Yeast Two-Hybrid Screening

Yeast two-hybrid (Y2H) screening was performed by Hybrigenics Services (www.hybrigenics-services.com) using a LexA-based system to identify protein-protein interactions. A bait construct encoding *C. elegans* PGL-1 (aa 450-672) was cloned into a vector for expression in yeast and screened against *C. elegans* Mixed Stages cDNA library fused to a Gal4 activation domain. The screen was conducted under selective conditions using 3-amino-1,2,4-triazole (3-AT) to suppress background growth from leaky HIS3 reporter expression. 83 million clones (10-fold the complexity of the library) were screened using a mating approach as previously described [63]. 260 His+ colonies were selected on a medium lacking tryptophan, leucine and histidine. The prey fragments of the positive clones were amplified by PCR and sequenced at their 5’ and 3’ junctions. Each identified prey clone was assigned a Predicted Biological Score (PBS, [64–66]) a statistical confidence score reflecting the likelihood of a true positive interaction. PBS scores ranged from A (highest confidence) to F (experimentally flagged artifact), based on the number and independence of prey fragments recovered, as well as empirical data from Hybrigenics’ extensive internal interaction database (>9,000 screens). Interactions flagged as PBS E represent prey domains with high connectivity across unrelated baits, indicating potential biochemical promiscuity or common protein interaction motifs. Prey fragments were mapped and analyzed using the DomSight tool, which compares the selected interacting domains (SIDs) to annotated functional and structural domains. Only interactions with high PBS scores were considered for further analysis (**Supplemental File 1**).

Further description of the confidence score:

The PBS relies on two different levels of analysis. Firstly, a local score takes into account the redundancy and independency of prey fragments, as well as the distribution of reading frames and stop codons in overlapping fragments. Secondly, a global score takes into account the interactions found in all the screens performed at Hybrigenics using the same library. This global score represents the probability of an interaction being nonspecific. For practical use, the scores were divided into four categories, from A (highest confidence) to D (lowest confidence). A fifth category (E) specifically flags interactions involving highly connected prey domains previously found several times in screens performed on libraries derived from the same organism. Finally, several of these highly connected domains have been confirmed as false-positives of the technique and are now tagged as F. The PBS scores have been shown to positively correlate with the biological significance of interactions [64, 66].

### eCLIP library preparation and sequencing

Enhanced crosslink and immunoprecipitation (eCLIP) experiments were performed according to the published protocol [67], except that dsRNA ligation was not performed as IFE-1 is a single-stranded RNA binding protein and that 250 U of RNase I (Ambion) was treated at 37°C for 10 minutes with constant shaking at 1200 rpm prior to immunoprecipitation. Briefly, upon RNase I digestion, 2% of 5 mg worm lysate was taken as inputs and the rest of the lysate was incubated with 100 µl of anti-FLAG magnetic beads (Sigma) with a final concentration of 1:5 dilution of wash buffer (20 mM Tris-HCl (pH 7.5), 10 mM MgCl_2_, 5 mM NaCl, 0.2% Tween-20) for 1 hour. IP samples were washed once with high-salt wash buffer (50 mM Tris-HCl (pH 7.5), 1 M NaCl, 1 mM EDTA, 0.1% SDS, 1% NP-40) and washed twice with the wash buffer. T4 PNK Minus reaction was performed on bead using T4 PNK Minus (NEB) at 37°C for 20 minutes with constant shaking at 1200 rpm, followed by 1x wash with high-salt wash buffer and 3x wash with wash buffer. IP samples were subjected to Fast AP (Thermo Scientific) reaction at 37°C for 10 minutes with constant shaking at 1200 rpm and PNK reaction with T4 PNK (NEB) at 37 °C for 20 minutes with constant shaking at 1200 rpm without removing the FastAP reaction mixture. Then, IP samples were washed 1x with high-salt wash buffer and 3x with wash buffer. 3’ RNA adapter (CLASHInvRiL19, **Table S1**) was ligated on bead using T4 RNA ligase (high-concentration, NEB) at RT for 75 minutes on a rotator. IP samples were then washed 1x with wash buffer, 1x with high-salt wash buffer, and 2x with wash buffer. All input and IP samples were denatured using 4xNuPAGE™ sample buffer (Invitrogen) at 65°C for 10 minutes and resolved in NuPAGE™ 4-12% Bis-Tris gel (Invitrogen) at 150 V for 75 minutes, followed by transfer to nitrocellulose membrane at 30 V for overnight. After confirming the position of IFE-1 protein by western blot using FLAG antibody (Sigma), regions above IFE-1 protein (up to ∼250 kDa) were excised for proteinase K (NEB) treatment at 37°C for 40 minutes with constant shaking at 1200 rpm. RNA was isolated from proteinase K treatment reaction using Zymo RNA clean & concentrator kit (Zymo Research). Upon RNA isolation, input samples were also treated with FastAP (Thermo Scientific) for 10 minutes and T4 PNK (NEB) for 20 minutes, both at 37 °C with constant shaking at 1200 rpm, followed by clean up using Zymo RNA clean & concentrator kit (Zymo Research). 3’ RNA adapter (CLASHInvRiL19, **Table S1**) was ligated to input samples after incubation at 65°C for 2 minutes using T4 RNA ligase (high-concentration, NEB) at RT for 60 minutes with flicking every 15 minutes. Input samples were cleaned up using MyONE Silane beads (Invitrogen). Reverse transcription was performed for both IP and input samples after incubation at 65°C for 2 minutes using an RT primer (InvAR17, **Table S1**) and then at 55 °C for 20 minutes using Superscript IV (Invitrogen). The reaction was cleaned up using ExoSAP-IT (Applied Biosystems) and MyONE Silane beads (Invitrogen). Then, 5’ DNA adapter (Invrand10_3Tr3, **Table S1**) was ligated after incubation at 70°C for 2 minutes and then at RT for overnight using T4 RNA ligase (high-concentration, NEB) and 5’ deadenylase (NEB). Samples were cleaned up using MyONE Silane beads (Invitrogen) and PCR amplified for 8 cycles (Input) and 13 cycles (IP) using Q5 High-Fidelity PCR master mix (NEB) and PCR forward and reverse primers with barcodes (**Table S1**). Libraries were purified using Ampure XP beads (Beckman Coulter) and quantified using the D1000 high sensitivity screen tape system (Agilent, La Jolla, CA) before pooling. Pooled library was sequenced on NextSeq2000 (Illumina, San Diego, CA) at center of bioinformatics and genomics (CGB) at Indiana University Bloomington using paired-end sequencing (2X101 bp). RNA visualization of eCLIP samples were performed as previously described [67]. 5% of IP samples were subjected to SDS-PAGE and transferred to nitrocellulose membrane. Detection of biotin-labeled RNA was performed using a Chemiluminescent Nucleic Acid Detection Module Kit (Thermo Scientific). Immunoblot of eCLIP samples were performed with 2% of input samples and 2.5% of IP samples using antibodies against FLAG (Sigma). The immunoblot images without saturation were acquired using Image Lab software (version 6.1.0 build 7) on the Bio-Rad ChemiDoc imaging system.

### CLIP Sequencing Analysis

Fastq raw sequencing files were assessed for quality and filtered to remove adapters by fastp (v0.21.0) [68]. Barcodes were subsequently removed and reads deduplicated. Paired reads were merged with NGmerge (v0.03) [69]. Processed reads were mapped to the *C. elegans* genome using STAR (v2.7.10b) [70]. Custom perl scripts were used to identify RNA start sites and read coverage. Targets were identified using the CLIPPER software package [71]. For the 5’ ratio analysis, two ratio statistics were computed for each transcript in the *C. elegans* genome by dividing the reads in the IP samples by the paired input sample for the first 20% of the transcript (5’ end) and the remaining 80% of the transcript body. The ratio statistics were then compared, using the coverage in the body to normalize for the relative depth of sequencing present in the sample to identify mRNAs that have a high ratio of 5’ reads. RNAs less than 300 base pairs, along with RNAs that had a ratio of ratios statistic of 1:1 were removed to minimize background and false positives. The ratio of ratios was calculated independently for both biological replicates and then averaged. Transcripts were then ranked by their ratio of ratios statistic, so that transcripts with high proportion of 5’ enrichment would emerge at the top of the list. Tissue enrichment and gene ontology analyses were performed using the Wormbase tool [72]. RNA type and chromosome distribution pie charts were generated in the R software package (v4.5.0).

### RNA visualization of IFE-1-bound RNAs

RNA visualization of eCLIP samples were performed as previously described [67]. 5% of IP samples were subjected to SDS-PAGE and transferred to nitrocellulose membrane. Detection of biotin-labeled RNA was performed using a Chemiluminescent Nucleic Acid Detection Module Kit (Thermo Scientific).

### Immunoblot of input and CLIP samples

Western blot of eCLIP samples were performed with 2% of input samples and 2.5% of IP samples using antibodies against FLAG (Sigma). The western blot images without saturation were acquired using Image Lab software (version 6.1.0 build 7) in the BIO-RAD ChemiDoc imaging system.

### Protein expression and purification

*E. coli* codon optimized sequences of six histidine, TEV cleavage site tagged IFE-1 and PGL-1 were commercially designed and ordered (Twist Biosciences). Halo and six histidine tagged PGL-3 was cloned from N2 wildtype and HaloTag strain cDNA with superscript IV (Invitrogen). These amplified constructs were inserted in a pET21a(+) protein expression vector (Novagen) with Gibson cloning [73]. Sanger sequencing confirmed proper construct design and insertion.

The protein expression plasmids were transformed into LOBSTR cells (Kerafast) and selected on Ampicillin/Chloramphenicol Luria Broth (LB) plates. Small LB cultures were grown in Ampicillin (50 µg/ml) and Chloramphenicol (25 µg/ml) overnight at 37°C with shaking (225 rpm). 1L LB cultures with Ampicillin (50 µg/ml) and Chloramphenicol (25 µg/ml) were grown at 37°C with shaking (225 rpm) until ∼0.8 OD (A600), cooled for 30–120 min, and induced with a final concentration of 0.1 mM IPTG. Cultures were then grown at 16°C with shaking (160 rpm) for 16–20 h, collected, frozen in liquid nitrogen, and stored at -80°C until use. Bacterial pellets were defrosted on ice and reconstituted in lysis buffer (20 mM Tris-HCl pH 8.0, 1M NaCl, 10 mM imidazole, 5 mM beta-mercaptoethanol (BME)) with protease inhibitors (cOmplete™ EDTA-free, Roche, Indianapolis, IN). Samples were lysed with a microfluidizer [74]. Lysed samples were centrifuged at low (3220 x *g*, 4 °C, 30 min), and the supernatant incubated with 1 ml NiNTA beads (Thermo Fisher Scientific, Waltham, MA) for 2 hours at 4 °C with rotation. Sample supernatant was removed by gravity flow on a column, washed with 50 ml lysis buffer, and eluted using lysis buffer with increasing imidazole concentrations (20, 40, 60, 80, 100, 250 mM). Eluted samples were monitored via Bradford assay (Bio-Rad, Hercules, CA), and samples containing recombinant protein were dialyzed overnight in HN buffer (IFE-1: 20 mM HEPES pH 7, 100 mM NaCl, 1 mM DTT; PGL: 20 mM HEPES pH 7, 500 mM NaCl, 1 mM DTT). The dialyzed samples were concentrated with a Vivaspin 3K or 10K concentrator (Sartorius, Göttingen, Germany) for IFE-1 and PGL, respectively. SDS-PAGE gels and Coomassie staining were used to assess protein purity. Purified protein samples were frozen in liquid nitrogen and stored at -80°C until use.

### Protein binding assay

Purified His-tagged IFE-1 and PGL proteins were incubated with 7-methyl-guanosine-5’-triphosphate (m^7^GTP)-agarose beads (Jena Bioscience and Creative BioMart) for 2 hours at 4 °C with gentle rotation. Following incubation, the mixtures were transferred to spin columns (Thermo Scientific) and washed five times with wash buffer (20 mM HEPES pH 7.5, 150 mM NaCl, 1 mM EDTA, 1 mM DTT, 0.1% Tween-20) by centrifugation at 2000 x g for 1 minute at 4 °C. Bound proteins were eluted in 5X SDS sample buffer and analyzed by SDS-PAGE followed by Coomassie staining. Stained gels were imaged on a Bio-Rad ChemiDoc imaging system.

### Statistics

All statistical tests were performed using GraphPad Prism v.10.2.3 software. For comparison of two groups, significance was determined using an unpaired *t*-test. For comparison of three or more groups, significance was determined using a one-way ANOVA. In all figures, error bars represent 95% confidence intervals (CI). ‘ns’ indicates not significant; \**P*<0.05; \*\**P*<0.01; \*\*\**P*<0.001; \*\*\*\**P*<0.0001.

### Alphafold3 modeling

The described protein sequences were submitted to the Alphafold3 [46] server, and the pTM and ipTM values for the generated model reported. Structure images were depicted by PyMOL (The PyMOL Molecular Graphics System, Version 3.0 Schrödinger, LLC).

### Data Availability

The *C. elegans* strains generated in this study have been deposited at the Caenorhabditis Genetics Center (CGC; https://cgc.umn.edu) or are available upon request. All plasmids are available upon request. All raw ChIP-seq sequencing data are available through the Gene Expression omnibus (GEO; Entry ID: GSE299469).

## Supplemental Figure captions

**Figure S1.**
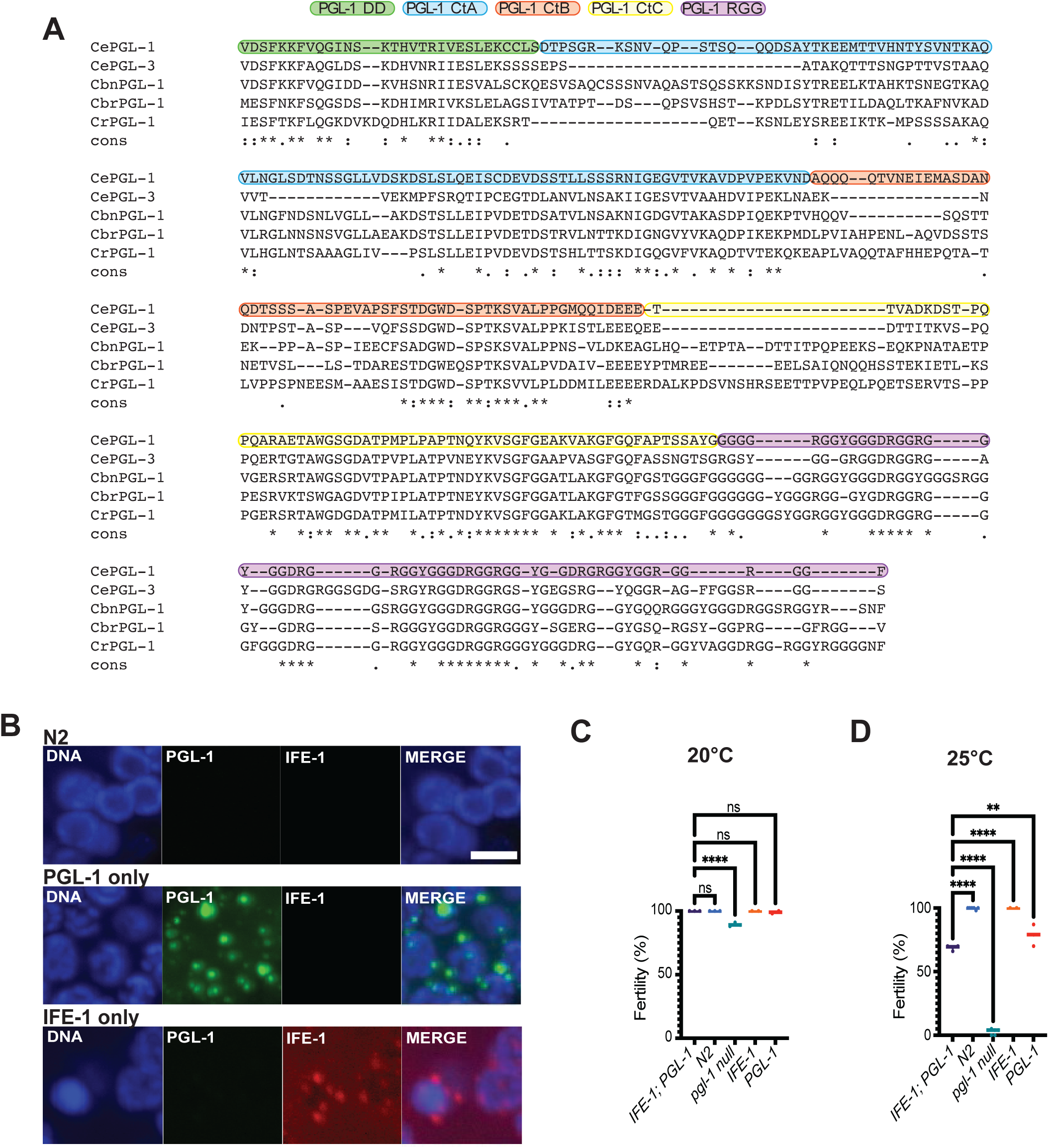
PGL-1 sequence alignments and additional imaging and fertility results. **(A)** Sequence alignment of PGL-1 and PGL-3 dimerization domain (DD; green), CtA (light blue), CtB (orange), CtC (yellow), and RGG (purple) regions in *C. elegans* (Ce), *C. brenneri* (Cbn), *C. briggsae* (Cbr), *C. remanei* (Cr). Alignment and conservation (cons.) determined by T-Coffee [75]. Starred (*) residue alignments are identical, while period (.) and colon (:) residues are similar. (**B**) Confocal microscopy images of adult N2 wildtype, PGL-1::V5::Halo (PGL-1), and IFE-1::3xFLAG::mScarlet (IFE-1) worms. DNA, DAPI, cyan; PGL-1, Halo Oregon Green, green; IFE-1, αFLAG, red; and a merged panel. All images represent 10-layer z-stack maximum intensity projections. White scale bar, 5 μm. (**C-D**) Fertility of N2 wildtype, *pgl-1(bn101) (pgl-1 null)*, PGL-1, IFE-1, and IFE-1; PGL-1 worms. Worms were plated individually, propagated at 20°C (C) or 25°C (D), and scored for presence of larval progeny after 4 days. Results reported as the percentage of fertile animals relative to the total number tested in each experiment. *n =* 3 independent experiments. ns, not significant; **, p-value <0.01; ****, p-value <0.0001.

**Figure S2.**
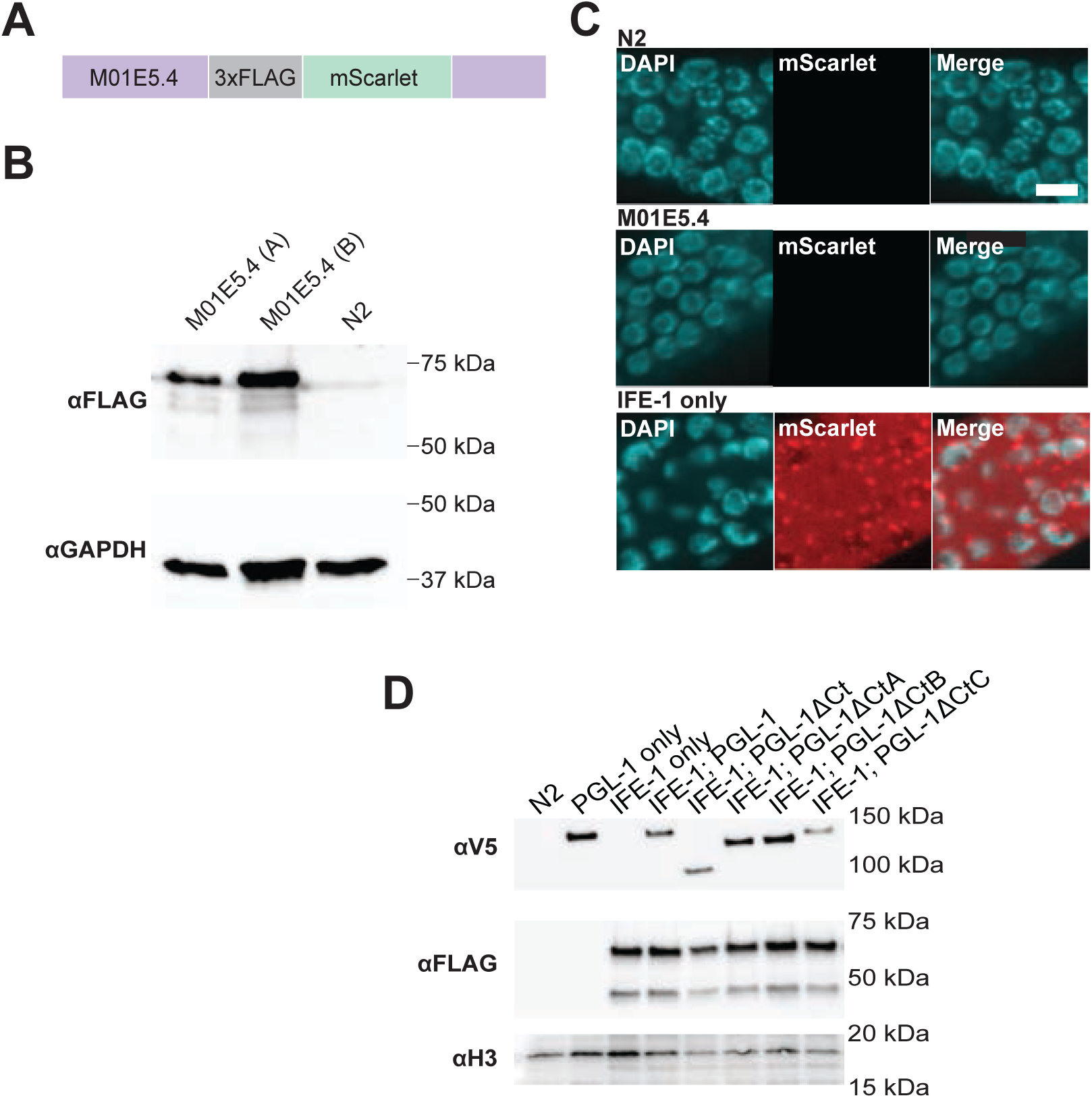
Analysis of M01E5.4 and protein expression of PGL-1 Ct mutants. **(A)** Linear diagram of *C. elegans M01E5.4* (purple) tagged with 3xFLAG (grey) and mScarlet (green). (**B**) Immunoblot of 3xFLAG::mScarlet tagged M01E5.4 (MO1E5.4) adult hermaphrodites. Two tagged M01E5.4 strains (“A” and “B”) and N2 wildtype worm samples were separated by SDS-PAGE and probed with FLAG and GAPDH antibodies. (**C**) Representative germline images of N2, M01E5.4, and IFE-1::3xFLAG::mScarlet (IFE-1) strains. DNA, DAPI, cyan; mScarlet, αFLAG, red; and a merged panel. All images represent 10-layer z-stack maximum intensity projections. Scale bar: 5 μm. (**D**) Immunoblot of adult hermaphrodite worm strains expressing IFE-1::3xFLAG::mScarlet (IFE-1) with PGL-1::V5::Halo (PGL-1), PGL-1ΔCt::V5::Halo (PGL-1ΔCt), PGL-1ΔCtA::V5::Halo (PGL-1ΔCtA), PGL-1ΔCtB::V5::Halo (PGL-1ΔCtB), and PGL-1ΔCtC::V5::Halo (PGL-1ΔCtC). N2 wildtype, PGL-1 only, IFE-1 only strains were used as controls. Samples were separated by SDS-PAGE and probed with V5, FLAG, and histone H3 (H3) antibodies.

**Figure S3.**
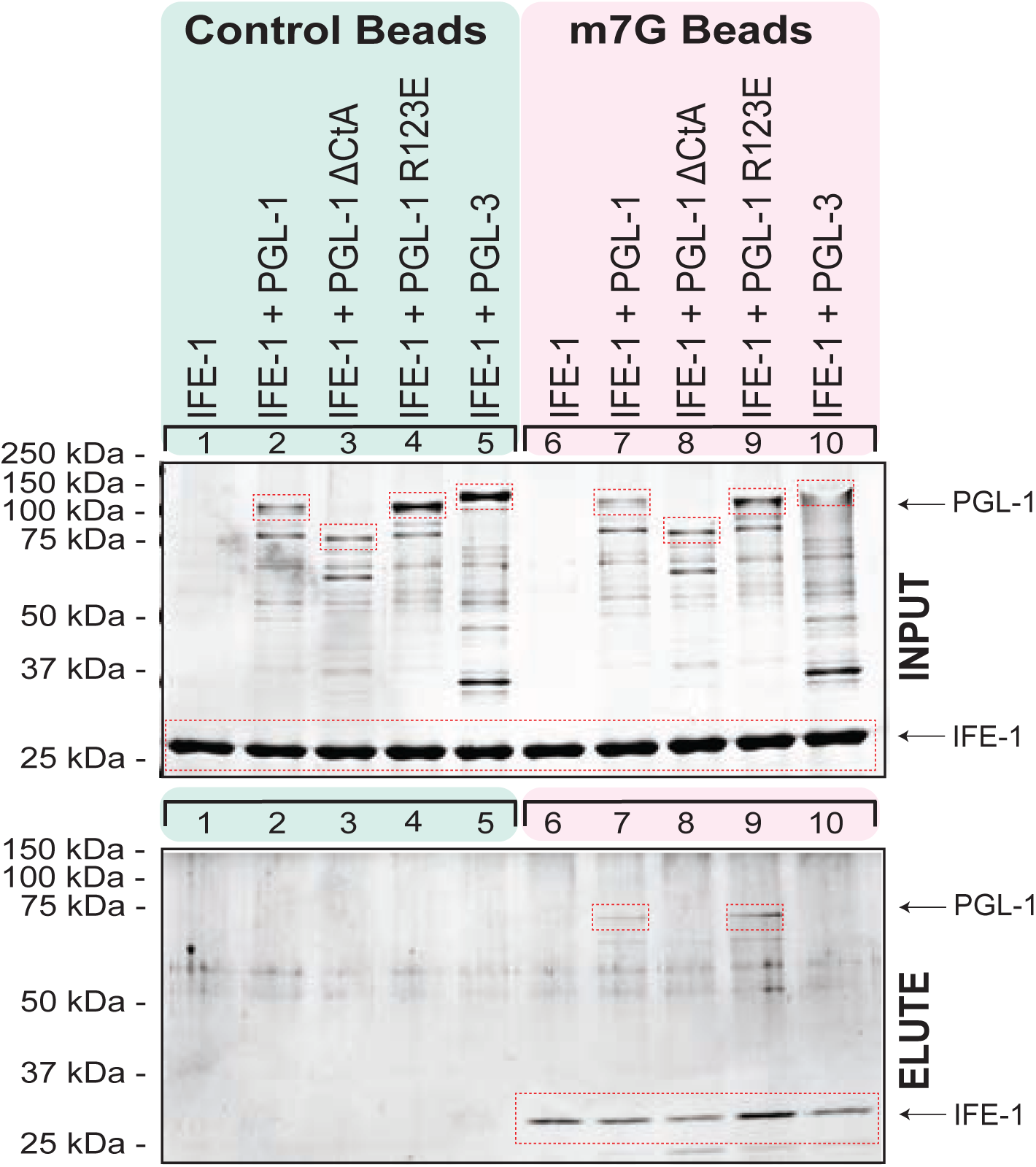
Recombinant purified IFE-1 binds directly to PGL-1 CtA *in vitro*. Recombinant IFE-1 was incubated with recombinant PGL-1, PGL-1ΔCtA, PGL-1 R123E, or PGL-3::Halo (PGL-3), and subjected to pulldown using either control beads or m^7^G agarose beads. Input and elution samples were analyzed by Coomassie-stained SDS-PAGE. Full length protein highlighted with red boxes. In the elution fraction, IFE-1 was retained on m^7^G beads, and co-precipitation was observed only with PGL-1 and PGL-1 R123E, indicating a specific interaction between IFE-1 and these PGL-1 variants.

**Figure S4.**
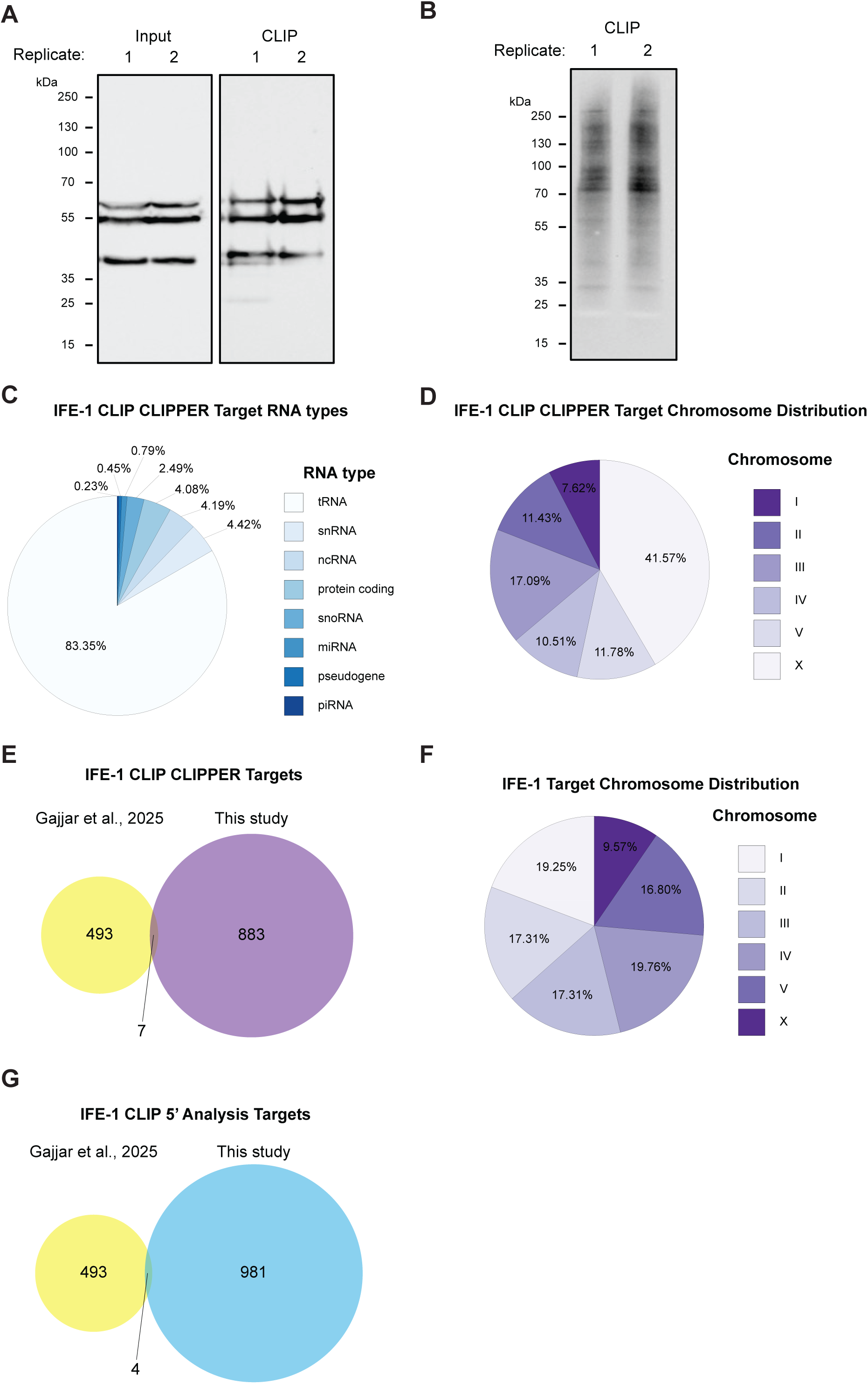
Additional eCLIP-seq results. **(A)** Immunoblot analysis of eCLIP input and IP samples, probed with a FLAG antibody. (**B**) Immunoblot of IFE-1 CLIP samples probed with streptavidin for biotin detection. Region above IFE-1::3xFLAG::mScarlet (∼70-250 kDa) was excised for downstream RNA isolation and sequencing. (**C**) RNA classification from the 883 RNAs identified as IFE-1 targets by CLIPPER analysis. (**D**) Chromosome distribution of 883 CLIPPER determined IFE-1 targets. (**E**) Overlap between 883 CLIPPER determined IFE-1 targets and 493 IFE-1 transcript targets identified previously [28]. (**F**) Overlap between the 982 top 25%, 5’ ratio analyzed IFE-1 mRNA transcripts and transcripts identified by [28]. (**G**) Chromosome distribution of IFE-1 associated mRNAs identified in 5’ ratio analysis.

**Figure S5.**
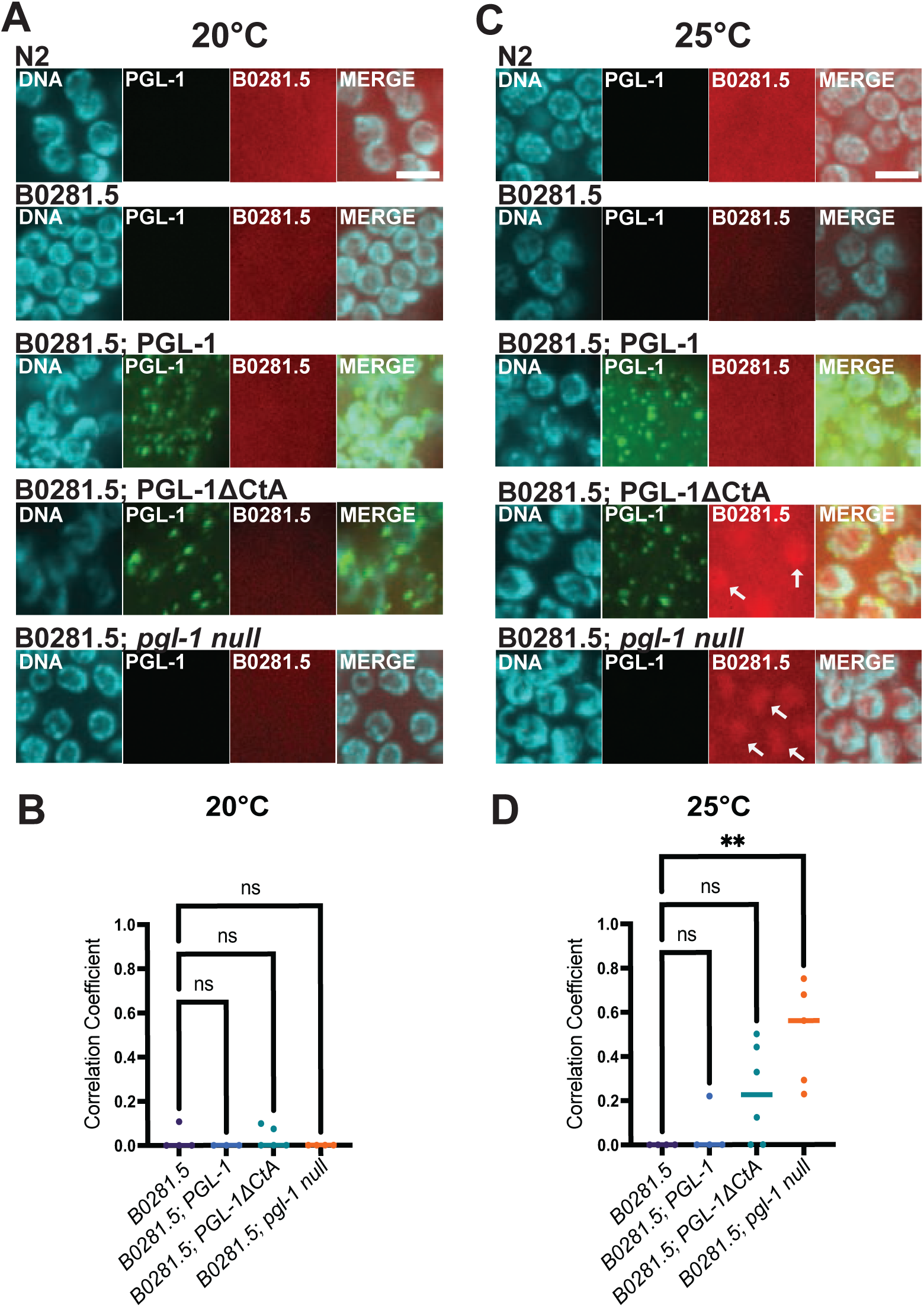
Germline B0281.5 protein expression is increased in *pgl-1 null (bn101)* background at elevated temperatures. Confocal microscopy images of adult hermaphrodite germlines. Strains expressed B0281.5::SNAP (B0281.5) with PGL-1::V5::Halo (PGL-1), PGL-1ΔCtA::V5::Halo (PGL-1ΔCtA), or *pgl-1*(*bn101*) (*pgl-1 null)*. Adult germlines were imaged by confocal microscopy. DNA, DAPI, cyan; PGL-1, Halo Oregon Green, green; B0281.5, SNAP TMR, red; and a merged panel. All images represent 10-layer z-stack maximum intensity projections. White scale bar, 5 μm. (**A**) Representative images of SNAP-tagged B0281.5 worms in PGL-1, PGL-1ΔCtA, and *pgl-1 null* genetic backgrounds, propagated at 20°C. (**B**) Quantification of B0281.5 signal colocalization with DNA at 20°C. (**C**) Representative images of the SNAP-tagged B0281.5 worms previously described, propagated at 25°C. White arrows highlight B0281.5 expression in PGL-1ΔCtA and *pgl-1 null* genetic backgrounds. Statistical significance was only observed in the *pgl-1 null* animals. (**D**) Quantification of B0281.5 colocalization with DNA at 25°C. ns, not significant; **, p-value <0.01.

**Figure S6.**
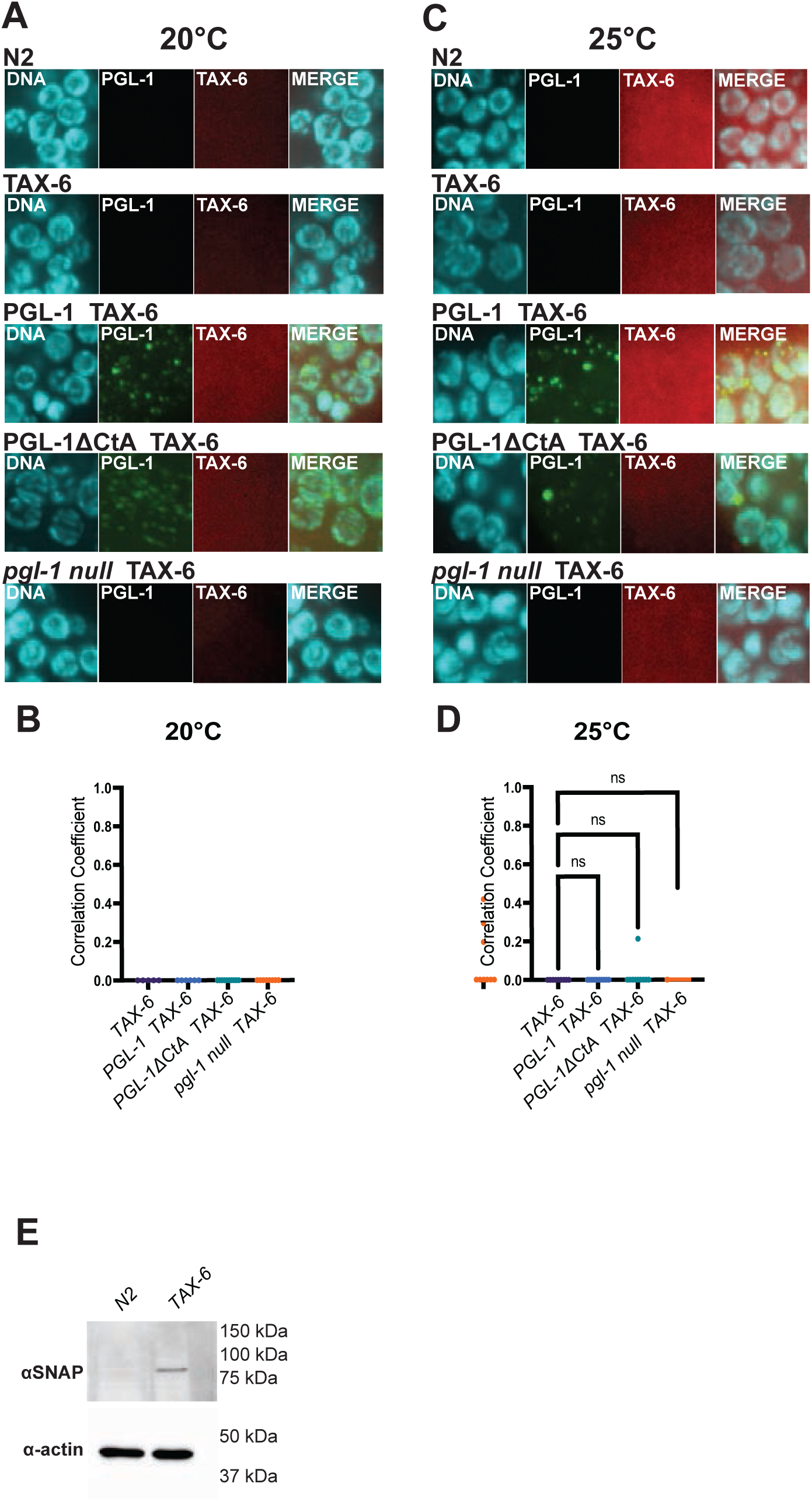
Germline TAX-6 protein expression is undetectable. Confocal microscopy images of adult hermaphrodite germlines. Strains expressed TAX-6::SNAP (TAX-6) with PGL-1::V5::Halo (PGL-1), PGL-1ΔCtA::V5::Halo (PGL-1ΔCtA), or *pgl-1*(*bn101*) (*pgl-1 null)*. Adult germlines were imaged by confocal microscopy. DNA, DAPI, cyan; PGL-1, Halo Oregon Green, green; TAX-6, SNAP TMR, red; and a merged panel. All images represent 10-layer z-stack maximum intensity projections. White scale bar, 5 μm. (**A**) Representative images of SNAP-tagged TAX-6 expression at 20°C in PGL-1, PGL-1ΔCtA, and *pgl-1 null* genetic backgrounds. (**B**) Quantification of TAX-6 colocalization with DNA at 20°C. (**C**) Representative images of SNAP-tagged TAX-6 mutants in PGL-1, PGL-1ΔCtA, and *pgl-1 null (bn101)* genetic backgrounds at 25°C. No TAX-6 expression was observed**. D)** Quantification of TAX-6 colocalization with DNA at 25°C. No TAX-6 expression was observed at either temperature. ns, not significant. (**E**) Immunoblot of N2 wildtype and TAX-6 worms. Worm samples were separated by SDS-PAGE and probed with SNAP and actin antibodies.

**Figure S7.**
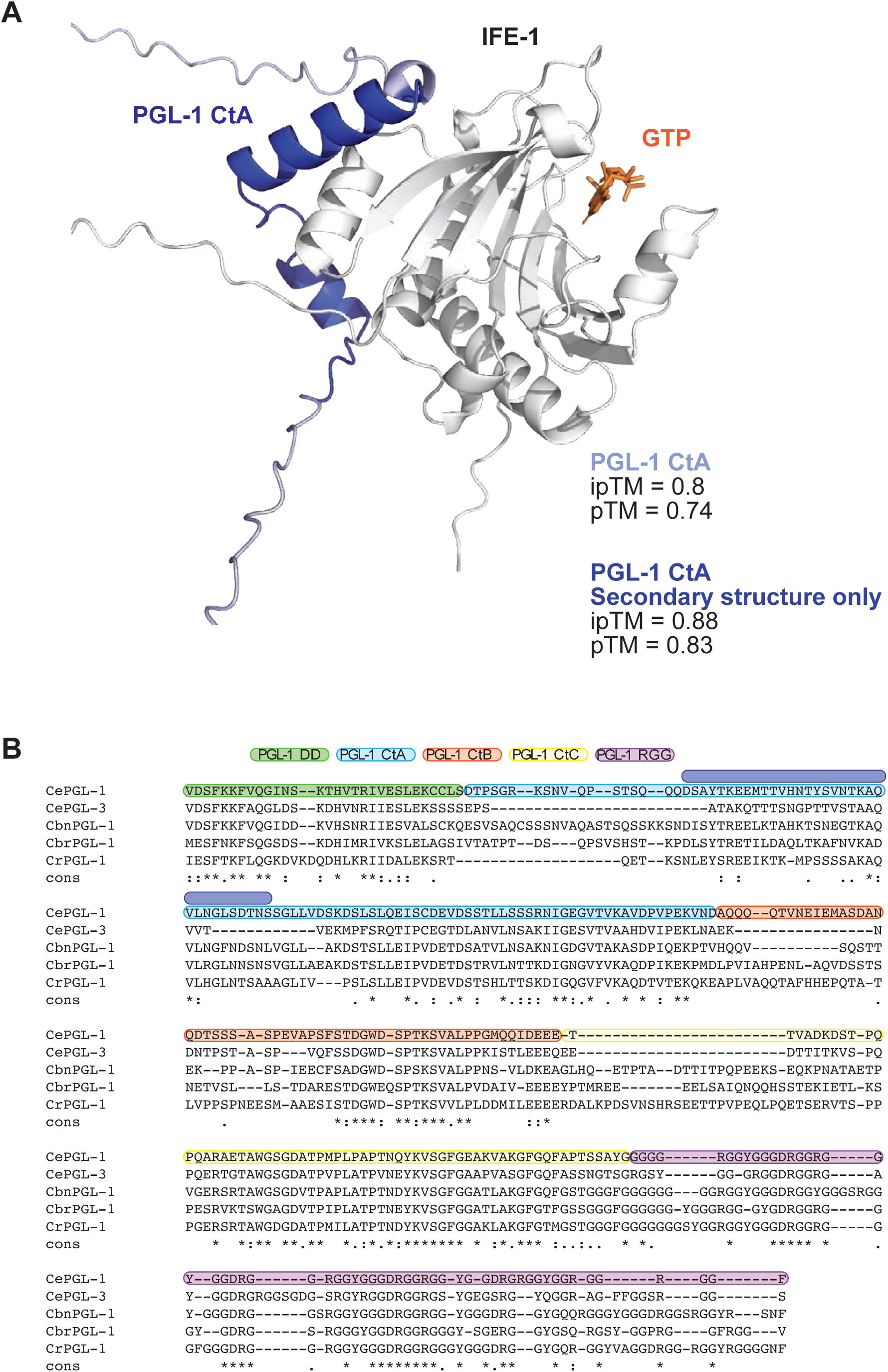
Alphafold3 prediction of the IFE-1 and PGL-1 CtA complex. **(A)** Alphafold3 [46] predicted structure of the PGL-1 CtA peptide bound to IFE-1. IFE-1, grey; PGL-1 CtA, blue; GTP, orange. ipTM and pTM values reported for the assemblies of PGL-1 CtA and PGL-1 CtA secondary structured region only. Images by PyMOL. (**B**) Sequence alignment of PGL-1 and PGL-3 dimerization domain (DD; green), CtA (light blue), PGL-1 CtA secondary structure (dark blue), CtB (orange), CtC (yellow), and RGG (purple) regions in *C. elegans* (Ce), *C. brenneri* (Cbn), *C. briggsae* (Cbr), *C. remanei* (Cr). Alignment and conservation (cons.) determined by T-Coffee [75]. Starred (*) residue alignments are identical, while period (.) and colon (:) residues are similar.

**Table S1.**
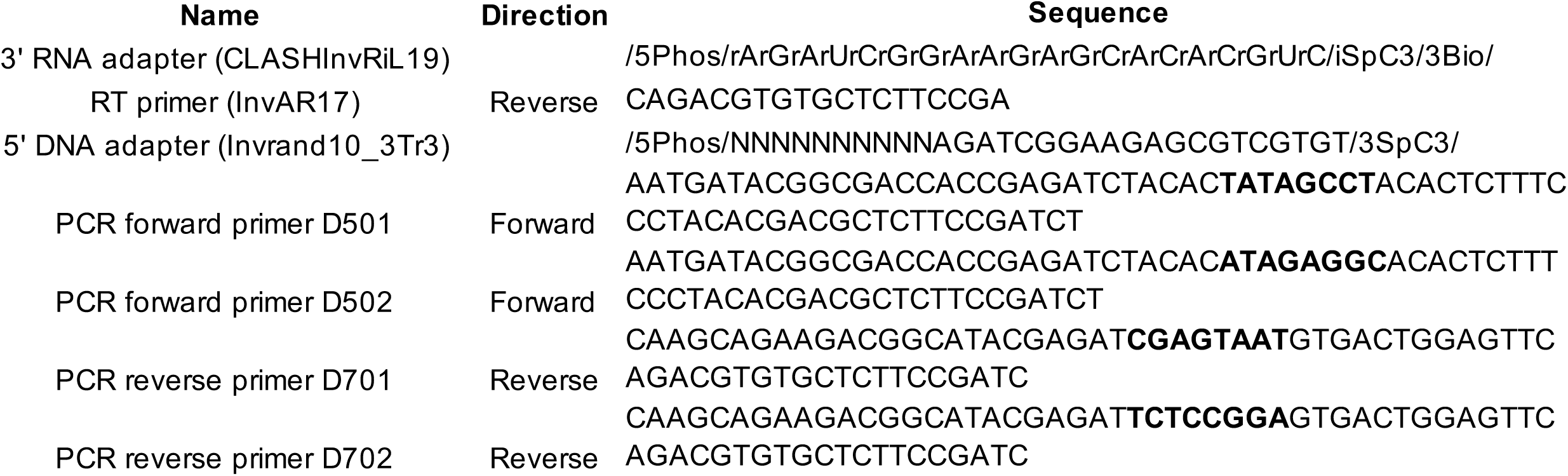
eCLIP oligonucleotide reagents.

**Supplemental File 1.**
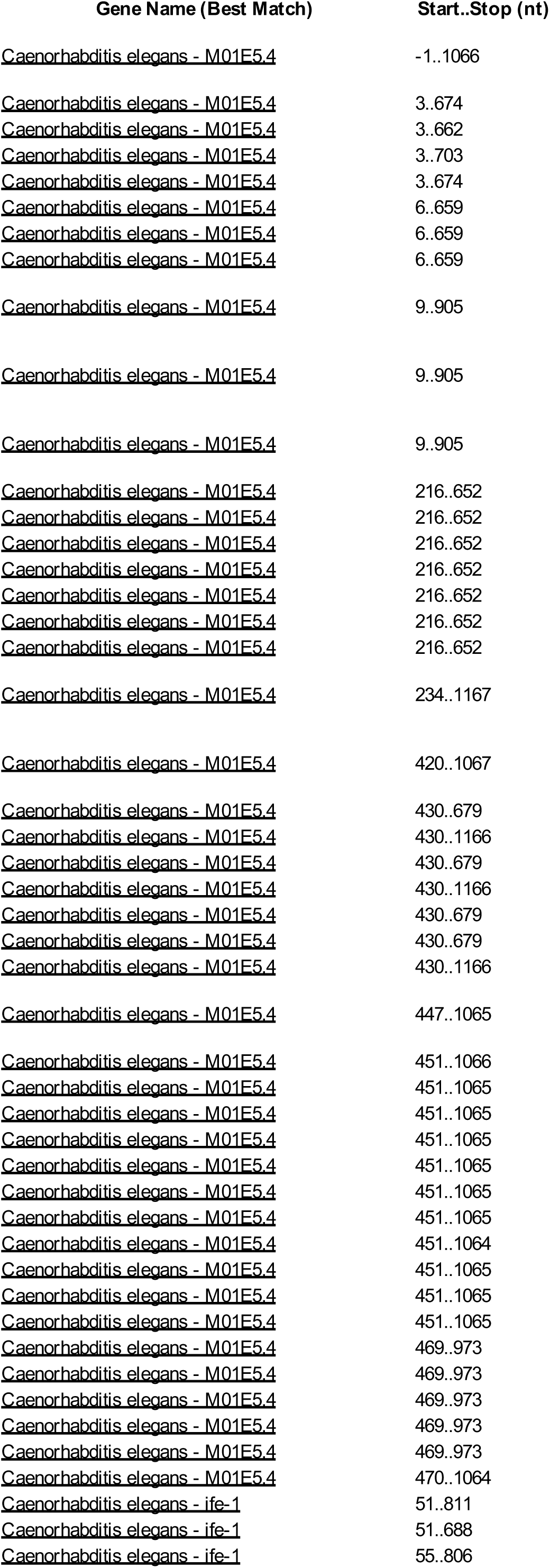
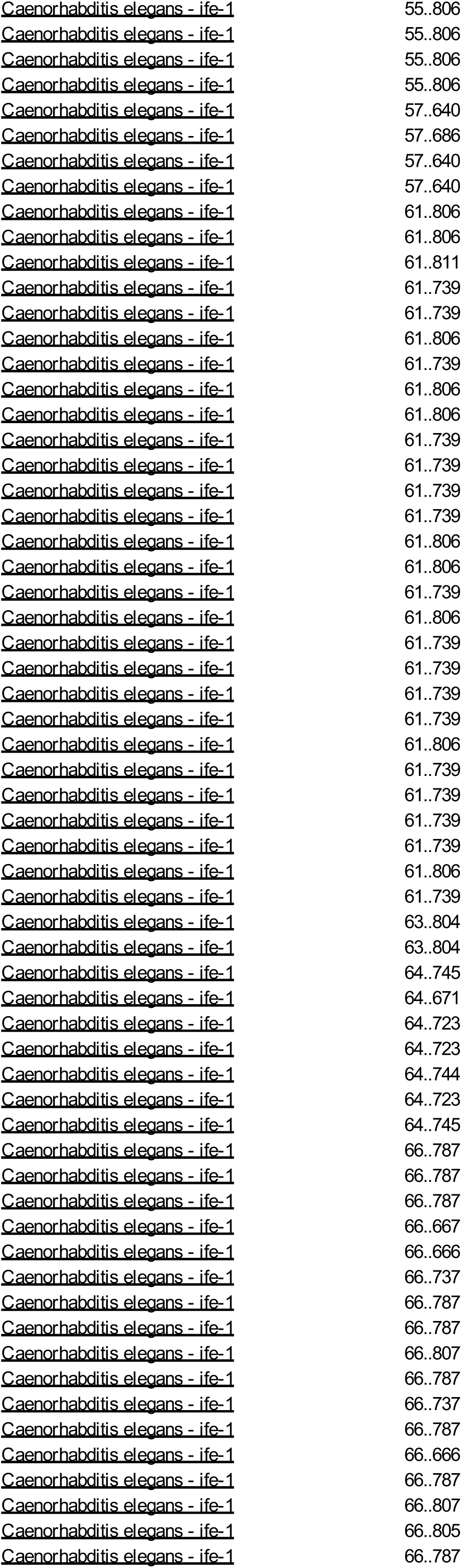
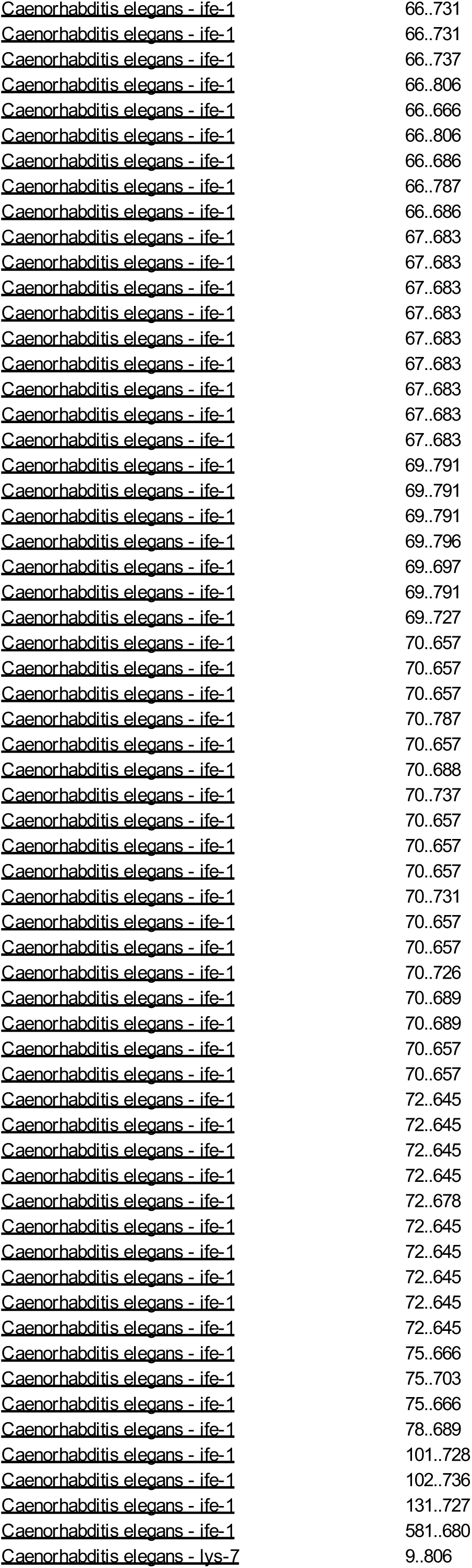
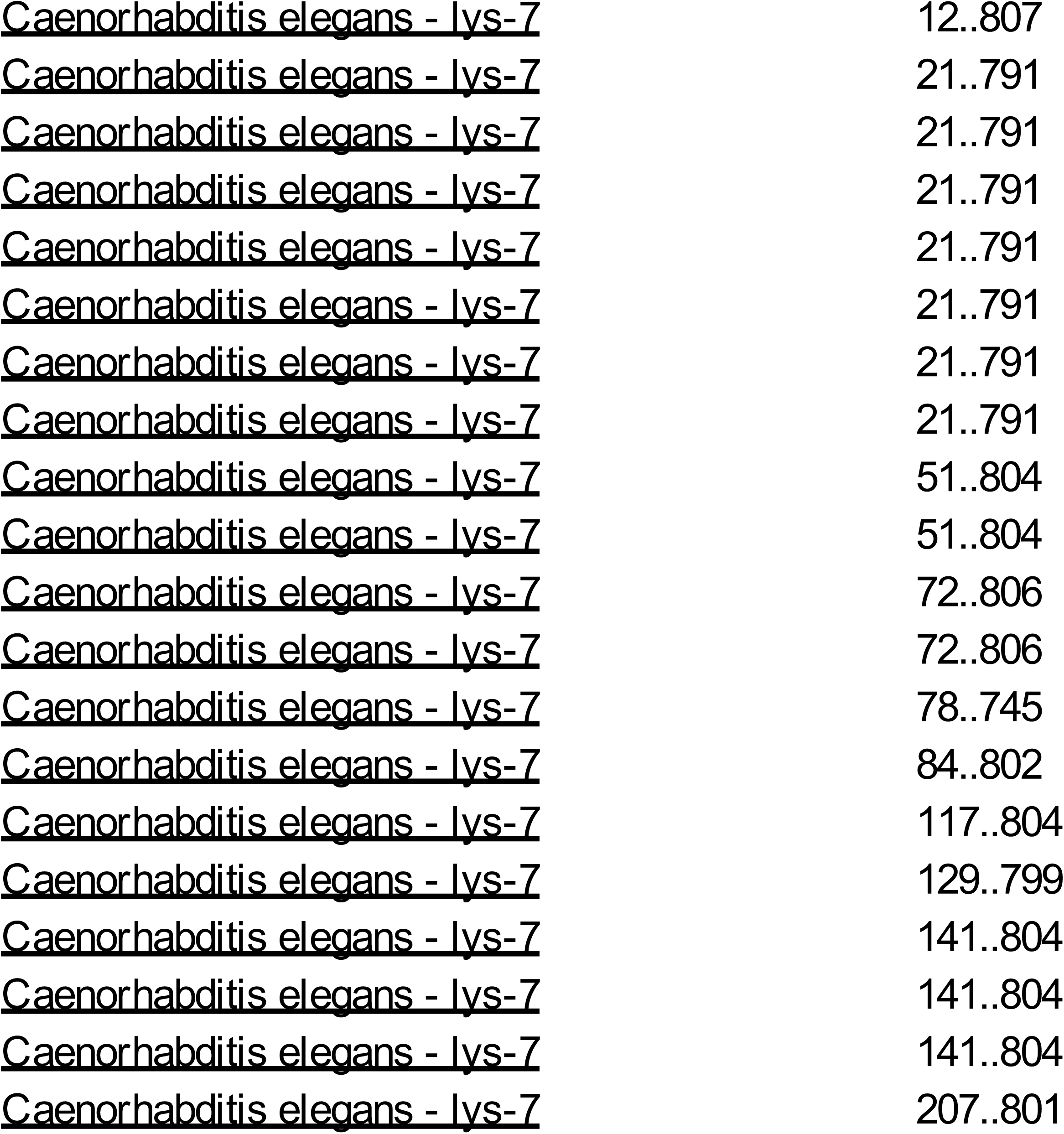
Yeast-two-hybrid interactors.

